# Variant to function mapping at single-cell resolution through network propagation

**DOI:** 10.1101/2022.01.23.477426

**Authors:** Fulong Yu, Liam D. Cato, Chen Weng, L. Alexander Liggett, Soyoung Jeon, Keren Xu, Charleston W.K. Chiang, Joseph L. Wiemels, Jonathan S. Weissman, Adam J. de Smith, Vijay G. Sankaran

## Abstract

With burgeoning human disease genetic associations and single-cell genomic atlases covering a range of tissues, there are unprecedented opportunities to systematically gain insights into the mechanisms of disease-causal variation. However, sparsity and noise, particularly in the context of single-cell epigenomic data, hamper the identification of disease- or trait-relevant cell types, states, and trajectories. To overcome these challenges, we have developed the SCAVENGE method, which maps causal variants to their relevant cellular context at single-cell resolution by employing the strategy of network propagation. We demonstrate how SCAVENGE can help identify key biological mechanisms underlying human genetic variation including enrichment of blood traits at distinct stages of human hematopoiesis, defining monocyte subsets that increase the risk for severe coronavirus disease 2019 (COVID-19), and identifying intermediate lymphocyte developmental states that are critical for predisposition to acute leukemia. Our approach not only provides a framework for enabling variant-to-function insights at single-cell resolution, but also suggests a more general strategy for maximizing the inferences that can be made using single-cell genomic data.

## INTRODUCTION

Our understanding of the genetic basis for a number of diseases has advanced considerably through the success of genome-wide association studies (GWAS)^1^. These studies have identified hundreds of thousands of variants associated with different complex phenotypes and have the potential to expand our knowledge of underlying biological mechanisms. Genetic fine-mapping has improved our ability to define putative causal variants^2^. However, the underlying mechanisms for the vast majority of variants remain undefined and this is particularly challenging given that most of these variants (~90%) appear to impact non-coding regulatory elements that are critical for gene expression^2,3^. While bespoke approaches have identified mechanisms at select loci, systematic functional mapping approaches could significantly accelerate our understanding of how genetic variation underlying complex diseases or phenotypes alters human biology.

Concomitantly, there have been enormous advances in the application of single-cell genomics to provide a refined and higher-resolution view of human biology than has been previously achievable^4^. This has motivated efforts to generate large single-cell genomic atlases^5,6,7,8^, including those measuring gene expression and/or epigenetic states, for a range of healthy and diseased human tissues. Such invaluable resources could enable efforts to map the functions of and tissue/cellular context for the enormous number of genetic variants that have been identified through GWAS. Co-localization of cell-type specific regulatory marks and genetic variants serves as a basis to perform such systematic functional inferences to identify relevant cellular contexts in which phenotype-associated variation is acting^9,10,11,12,13,14^. While these approaches have been applied to bulk-level or aggregated data, such methods would have further promise if they could be applied to robustly make inferences in single cells. However, this is unfortunately limited by the extensive amount of sparsity and noise (> 95% of sparsity, Supplementary Fig. 1a,b) found in single-cell epigenomic data^15^. The majority of cells are uninformative for most inferential approaches at specific loci, since the absence of signals may be attributable to technical or biological causes (Supplementary Fig. 2a).

This problem is analogous to early attempts to optimize performance of internet search engines, where variable and limited data contained on individual websites constrained the ability to rank the most relevant results. Google successfully addressed this issue through the development of the PageRank algorithm^16^ that is able to output the most important sites in a search in ranked order by generating a network of all websites and estimating the probability that an individual randomly clicking on links would arrive at a particular site in this network. This network propagation strategy has been successfully applied across a range of problems in biology, particularly in the context of noisy and incomplete observations^17^. Phenotypic characteristics and relatedness across individual cells can be well represented by high-dimensional features and distilled into a cell-to-cell network^18^. We reasoned that this network propagation strategy could enable efficient and improved assessment of phenotype-relevant cell types and states from single-cell genomic data with high sparsity and dropout (Supplementary Fig. 2b). Here, we have instantiated this network propagation approach through the SCAVENGE (**S**ingle **C**ell **A**nalysis of **V**ariant **E**nrichment through **N**etwork propagation of **GE**nomic data) method to optimize the inference of functional and genetic associations to specific cells at single-cell resolution. We discuss the development and validation of this approach, along with examples of the key biological insights that can be gained through application of this method. We provide new knowledge for how human genetic variation can alter distinct stages of hematopoiesis, how specific monocyte cell states can contribute to genetic risk for severe COVID-19, and how distinct intermediates in B cell development can underlie the predisposition to acquiring acute lymphoblastic leukemia - biological insights that would not have been possible without a robust method for mapping genetic variation to specific cell states at single-cell resolution.

## RESULTS

### Overview of SCAVENGE

Co-localization approaches using genetic variants and single-cell epigenomic data are unfortunately uninformative for many cells given the extensive sparsity across single-cell profiles. Therefore, only a few cells from the truly relevant population demonstrate relevance to a phenotype of interest. Nonetheless, the global high-dimensional features of individual single cells are sufficient to represent the underlying cell identities or states, which enables the relationships among such cells to be readily inferred^15^. By taking advantage of these attributes, SCAVENGE identifies the most phenotypically-enriched cells by co-localization and explores the transitive associations across the cell-to-cell network to assign each cell a probability representing the cell’s relevance to those phenotype-enriched cells via network propagation (Methods and Fig. 1).

For a disease or trait with fine-mapped causal variants and epigenomic information, such as the single-cell assay for transpose accessible chromatin through sequencing (scATAC-seq), we use g-chromVAR^13^ to calculate bias-corrected Z scores to estimate trait-relevance for each single cell (Fig. 1a) by integrating the probability of variant causality and quantitative strength of chromatin accessibility. Then, we rank the cells based on Z scores and use the cells with the top ranks to serve as seed cells, which are most relevant to the trait or disease of interest regardless of their identity. Since many cells may not show enrichment due to technical limitations (e.g. dropout), we then examine the relevance to the set of seed cells among the others by constructing a nearest-neighbor graph to capture the cell-to-cell topological structure in latent space (Methods and Supplementary Fig. 5a)^19,18,20^. In the nearest-neighbor graph, each node represents a cell and edges connect the most similar cells. We model the probability of relevance via a network propagation process, which can also be viewed as a Markov chain^21^, where the probability of a cell changes through a series of the random walk steps until the probability distribution converges at a stationary distribution (Methods and Fig. 1b). We project the seed cells on to the embedding graph and assign the seed cells to an even probability distribution representing the initial state of the cell-to-cell graph. As the graph is undirected, information propagated from the initial seed cells is only based on the structure and connectivity between cells in the graph. Once a stationary state is reached, SCAVENGE outputs the trait relevance score (TRS) for each single cell, which is a scaled and normalized probability distribution representing a quantitative metric for the relevance of a cell to a phenotype by accounting for both network structure and cell-to-cell distance (Methods). Thus, SCAVENGE’s TRS provides a unified measure of genetic trait/disease relevance that enables accurate annotation and characterization at the cell population and single-cell levels. From the TRS for the phenotype of interest, we could uncover previously unappreciated cell state/type associations among the continuum of states revealed through single-cell genomics (Fig. 1c).

**Fig.1:**
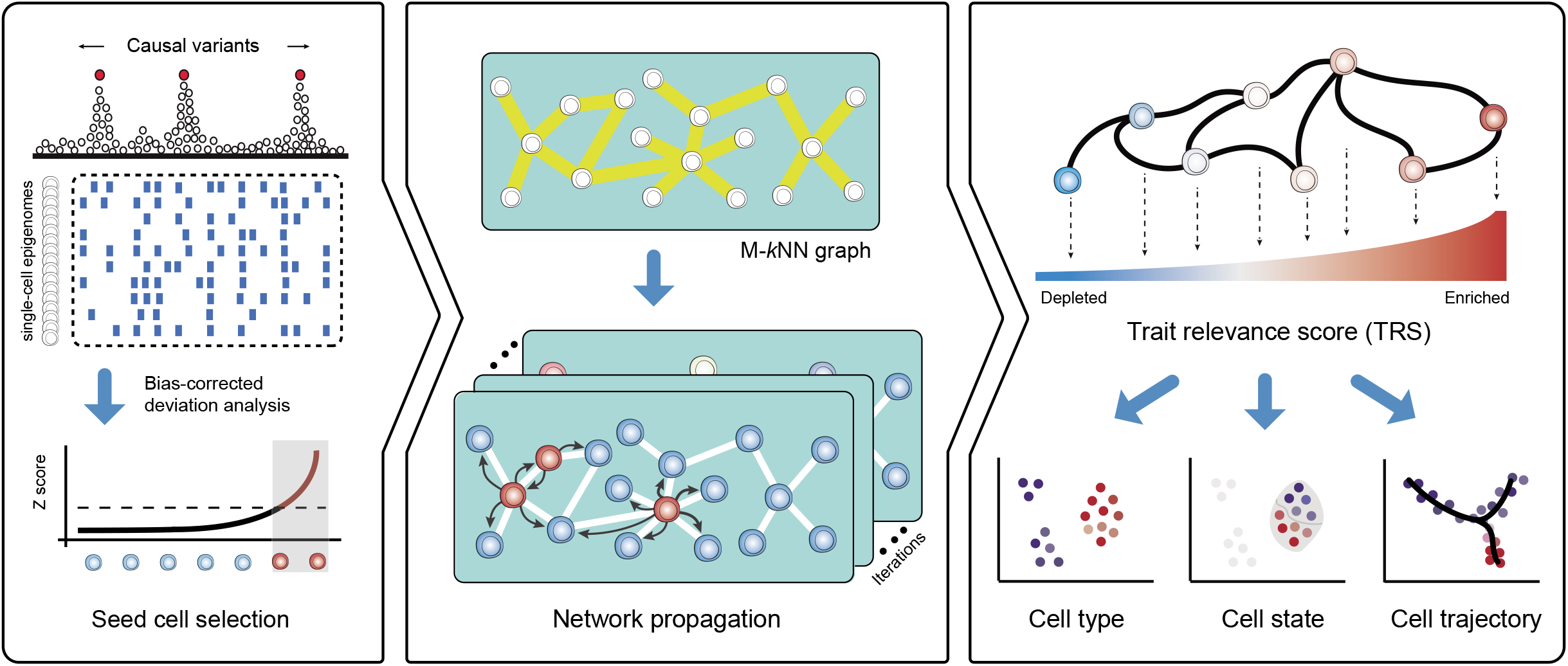
Overview of the SCAVENGE approach and application. **a**, For a given genetic trait/phenotype, the bias-corrected enrichment statistic is calculated for every single cell by integrating the posterior probabilities of fine-mapped variants and the scATAC-seq profiles. The top ranked cells are selected as the seed cells. **b**, A mutual *k* nearest neighbor (M-*k*NN) graph is constructed to represent cell–cell similarity, and the seed cells are projected on to this cell-to-cell graph. Network propagation scores for individual cells are defined according to the probability that the network reaches the stationary state from a number of steps of information propagation. **c**, Network propagation score is further scaled and normalized to obtain the per-cell SCAVENGE trait relevance score (TRS) that represents the relevance of trait/phenotype of interest for each single cell. Downstream analyses of functional annotation and interpretation are enabled at different levels including cell type, cell state, and cell trajectory.

### Benchmarking assessments with simulated and real data

To assess the power and accuracy of SCAVENGE, we first tested its performance on simulated scATAC-seq datasets (Methods). We employed genetic variants impacting monocyte count, which is a highly heritable phenotype^13,22^, as a test case. Relevant cell populations were characterized using bulk level ATAC-seq data (Supplementary Fig. 3) and we selected two cell types that were either enriched or depleted for relevance to this trait for an initial simulation involving the creation of synthetic single cell data. As expected, only a fraction of simulated cells presented accurate trait-associated relevance using traditional co-localization methods due to sparsity and technical noise, where those cells were mainly distributed in the top and bottom among the ranked group of cells (Fig. 2a). SCAVENGE considerably enhanced discovery of trait-relevant cells, with the improvement of the accuracy from 0.72 to 0.97, with particular improvements in accuracy for cells that previously showed moderate enrichment (Fig. 2a). This observation remained reproducible with the inclusion of simulated data from a larger array of cell types (Fig. 2b). These simulations also demonstrate that SCAVENGE is robust to a wide range of parameters, including the number of seed cells selected, number of neighbors used for graph construction, number of reads in baseline data, and variation in data noise levels (Supplementary Fig. 4a-d and Methods) To next assess whether SCAVENGE can detect this trait-relevant enrichment in real data, we applied SCAVENGE to a scATAC-seq dataset containing 4,562 peripheral blood mononuclear cells (PBMCs), of which the typical sparsity intrinsic to single-cell data has been demonstrated (Supplementary Fig. 1a,b, Supplementary Table 1). We found that SCAVENGE identified known trait-relevant cell populations with a high specificity (Fig. 2d,e,f, Supplementary Fig. 5b), which could not be achieved by randomly selected seed cells (Fig. 2g,h). In comparison, the enriched pattern was ambiguous and it was challenging to identify trait-relevant cells without SCAVENGE (Fig. 2c). These findings collectively show that SCAVENGE is robust and reproducible, which enables trait relevance to be correctly characterized at single-cell resolution.

**Fig.2:**
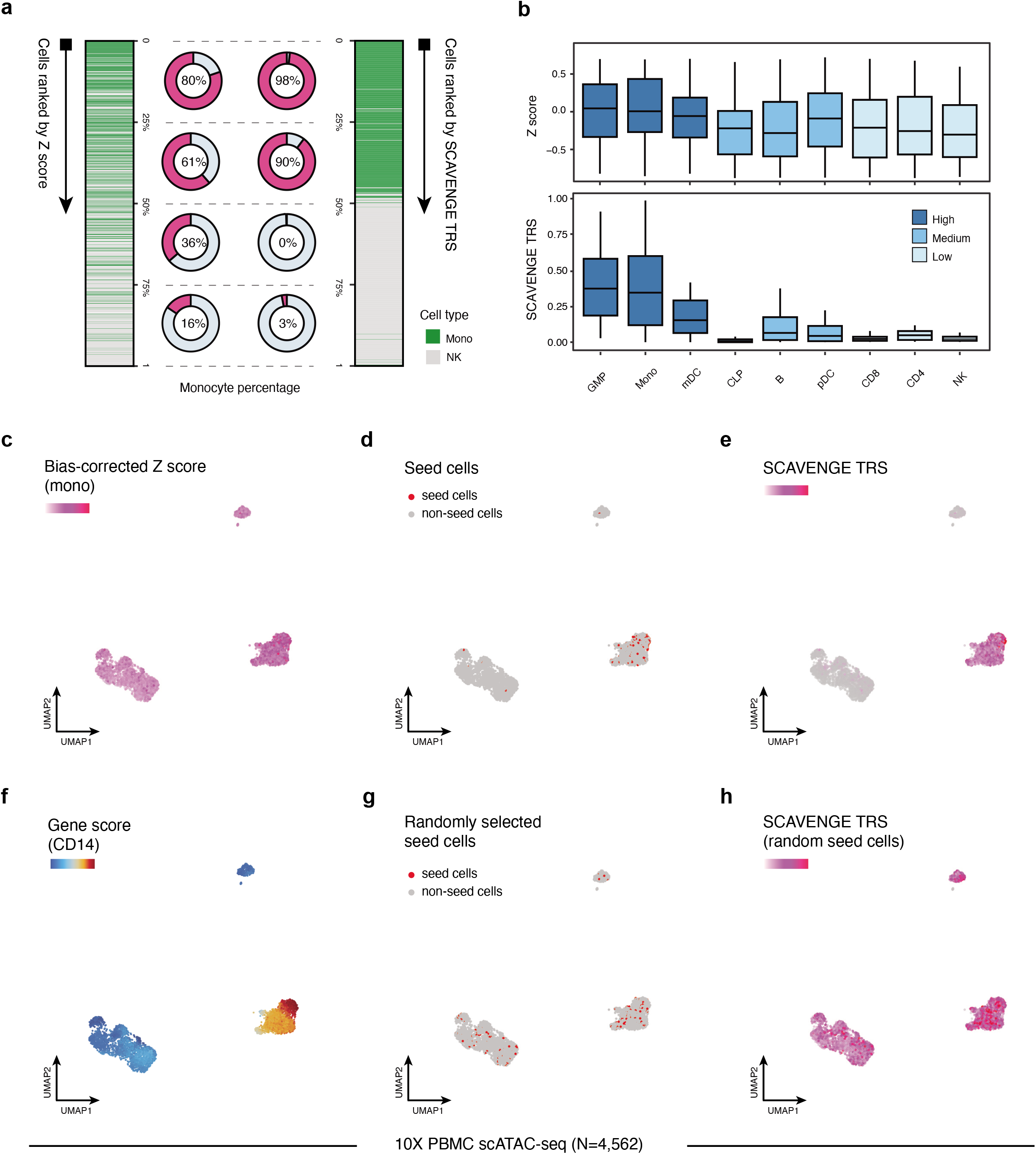
Assessing performance and robustness of SCAVENGE using simulated and real datasets. Two single-cell datasets were simulated from the same hematopoietic bulk ATAC-seq dataset (**a**), two cell types of monocyte (Mono) and natural killer cell (NK) are included, with 500 cells for each. (**b**), nine cell types of granulocyte-macrophage progenitors (GMP), Mono, myeloid dendritic cell (mDC), common lymphoid progenitor (CLP), B cell, plasmacytoid dendritic cells (pDC), CD8 T cell, CD4 T cell and NK are included, with 200 cells for each (Methods). The blood cell trait of monocyte count is investigated throughout simulated and real datasets. **a**, The cells are ranked according to the original bias-corrected Z score (left) and SCAVENGE network propagation score (right), respectively. The percentage of monocytes for each quarter is shown accordingly. **b**, The cells are ranked accordingly and the box plots depict the trait-relevant scores of cells from the second-quarter subsets before and after SCAVENGE. The box plot center line, limits and whiskers represent the median, quartiles and 1.5x interquartile range, respectively. **c-h**, Illustration of SCAVENGE analysis with a real hematopoietic scATAC-seq dataset. The UMAP embedding plots show (**c**) Z score, (**d**) seed cells, (**e**) SCAVENGE TRS obtained using seed cells in (**d)**, (**f**) gene accessibility score of canonical marker of monocyte, (**g**) randomly selected seed cells by matching the number of real ones, (**h**) SCAVENGE TRS obtained using seed cells in **g**. (**c, e**, and **h**) use the same color scheme, so that they can be compared across conditions.

### SCAVENGE enables systematic discovery of blood cell trait enrichments at distinct stages of human hematopoiesis

Following our initial benchmarking, we sought to assess whether SCAVENGE could accurately detect genetically-driven phenotype-relevant cell populations and generate biological insights with large-scale single-cell epigenomic data. We initially applied SCAVENGE to 33,819 scATAC-seq profiles covering the full spectrum of human hematopoietic differentiation from stem cells to their differentiated progeny^23^. We employed GWAS data from 22 highly heritable blood cell traits to examine causal cell states across this single-cell dataset^22^ (Supplementary Table 2). The TRSs of individual cells for four representative traits are shown in low-dimensional Uniform Manifold Approximation and Projection (UMAP) space (Fig. 3a-e). We found that traits related to relevant cell lineages showed distinct enrichments, illuminating cell-type specificity of these genetic effects that are well captured by SCAVENGE (Fig. 3a-e). For a single trait, the enriched cell compartments where the cells showed the highest TRS could be distributed far away from each other. For instance, two cell compartments enriched for eosinophil count were on opposite ends of the low-dimensional spatial projection (Fig. 3c). This suggests that network propagation will effectively capture trait-specific genetic associations in the relevant cell populations irrespective of proximity in the cell-to-cell graph. By aligning 23 hematopoietic cell populations previously annotated in bulk, we found the enriched cell compartments are highly concordant with our prior knowledge of causal cell types for blood traits^13,22,24^. For instance, lymphocyte count was most strongly enriched in CD4+ and CD8+ T cells, while mean reticulocyte volume was most strongly enriched in the late stages of erythroid differentiation (Fig. 3b,e).

**Fig.3:**
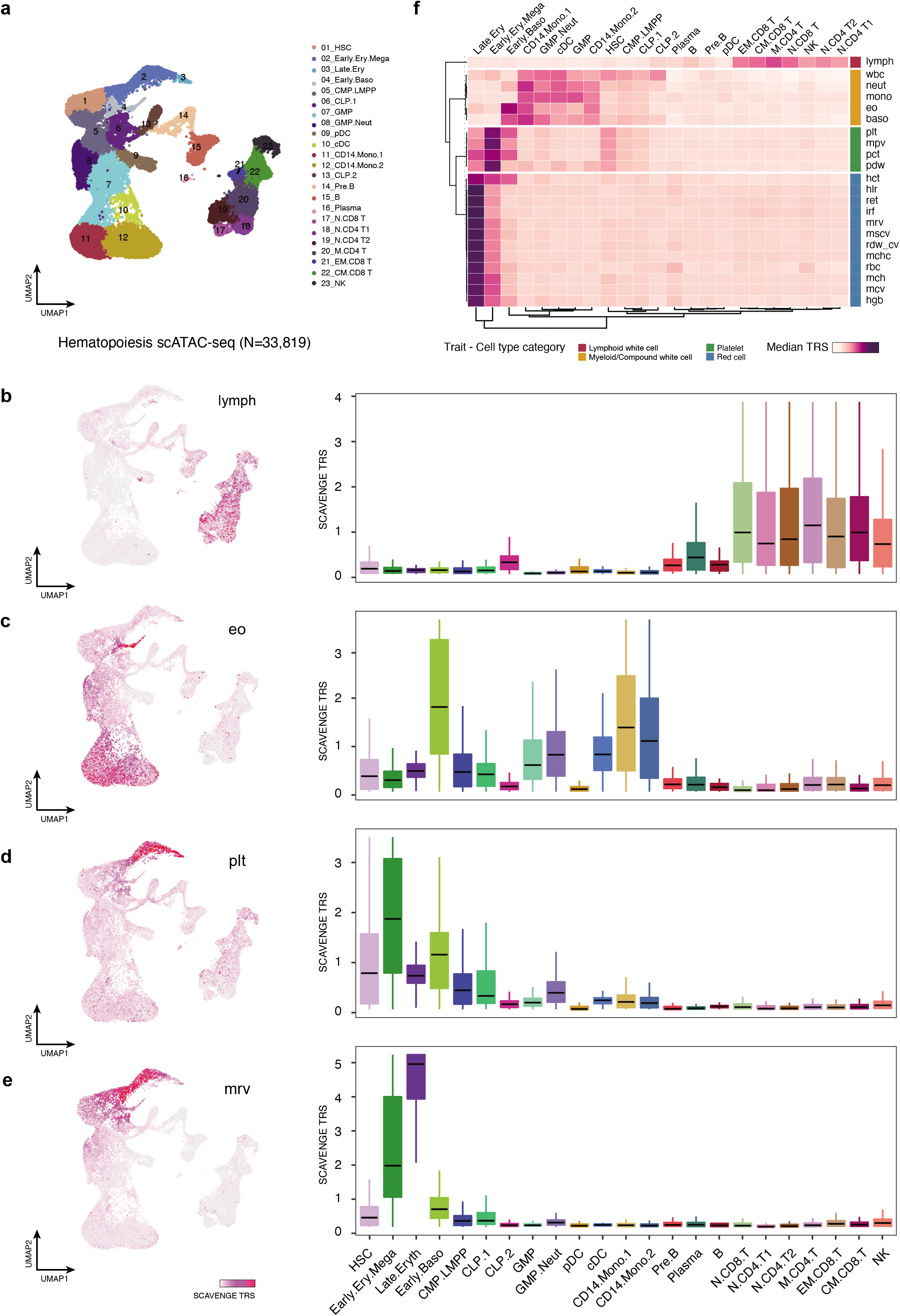
SCAVENGE enables comprehensive annotation of blood cell traits and captures the genetic basis of their causal cell types. The 22 blood cell traits are analyzed using SCAVENGE on a large hematopoiesis scATAC-seq dataset^23^. **a**, The UMAP plot shows the cell type labels. **b-e**, Per-cell SCAVENGE TRS for four representative traits including (**b**) lymphocyte count (lymph), (**c**) eosinophil count (eo), (**d**) platelet count (plt), and (**e**) mean reticulocyte volume (mrv) are shown in UMAP coordinates (left) and per cell type (right). Boxplots show the median with interquartile range (IQR) (25–75%); whiskers extend 1.5x the IQR. **f**, The median TRSs of cells belonging to the same cell type are shown in the heatmap. Unsupervised clustering analysis is performed and traits are grouped into four clusters using the *K*-means clustering algorithm. Trait-Cell type category is collected from a previous study^25^. HSC, hematopoietic stem cell; MPP, multipotent progenitor; CMP, common myeloid progenitors; LMPP, lymphoid-primed multipotent progenitor; MEP, megakaryocyte-erythroid progenitor; BMP, basophil-mast cell progenitor; N.CD4 T, naive CD4 T cell; N.CD8 T, naive CD8 T cell; M.CD4 T, memory CD4 T cell; CM.CD8 T, CD8 central memory T cell; CD8.EM T, CD8 effector memory T cell; Mega, megakaryocyte; Ery, erythrocyte; Baso, basophil; Mono, monocytes; Neut, neutrophil; cDC, classical dendritic cells. lymph, lymphocyte count; wbc, white blood count; neut, neutrophil count; mono, monocyte count; eo, eosinophil; baso, basophil count; plt, platelet count; mpv, mean platelet volume; pct, plateletcrit; pdw, platelet distribution width; hct, hematocrit; hlr, high light scatter reticulocyte percentage; ret, reticulocyte count; irf, immature fraction of reticulocytes; mrv, mean reticulocyte volume; mscv, mean sphered corpuscular volume; rdw_cv, red cell distribution width; mchc, mean corpuscular hemoglobin concentration; rbc, red blood cell count; mch, mean corpuscular hemoglobin; mcv, mean corpuscular volume; hgb, hemoglobin.

The single-cell profiling data enables more comprehensive cell profiling and better defined annotations than existing bulk-level data, providing a unique opportunity to explore previously undefined enrichments of specific cell types/states with particular genetic variants. To comprehensively explore genetic associations for hematological phenotypes in various cell contexts, we aggregated the SCAVENGE TRSs of cells within the same annotated cell type to define how specific hematopoietic traits are enriched at distinct stages of human hematopoiesis (Fig. 3f). Unsupervised clustering analysis demonstrated that different cell populations from the same lineage tend to have similar patterns across related and relevant traits. Reciprocally, traits relevant to the same hematopoietic lineage (e.g. red blood cell traits) were perfectly classified in the same module based on the cell-type TRS, consistent with our prior findings in bulk populations^25^, suggesting that SCAVENGE could provide insights into hematopoietic enrichments using single-cell data with unsupervised approaches as well as could be achieved with annotated bulk populations. Importantly, we found that SCAVENGE enables discoveries of novel cell-type enrichments and distinguishes enrichments for cell types with subtle differences, which are particularly challenging using existing bulk-level data. For example, basophil count is more strongly and specifically enriched in early basophils compared to other white blood cell traits. The complete white blood cell count is also enriched among hematopoietic stem cells (HSCs) and lymphoid progenitors in addition to different granulocyte progenitors and differentiated myeloid cells, consistent with contributions from various heterogeneous lineages to this phenotype. Red blood cell associated traits are more strongly enriched in differentiated erythroid cells than progenitors, while platelet associated traits tend to be enriched in early hematopoietic stem and progenitor populations, consistent with the earlier differentiation and commitment to the megakaryocytic lineage^26,27^.

To assess the validity and reproducibility of these findings, we also applied SCAVENGE to an independent scATAC-seq dataset with comprehensive coverage of human hematopoiesis that contained 63,882 cells with a more diverse cell-type composition^28^ (Supplementary Table 3). The results are consistent with our initial analysis and genetic correlations across the full spectrum of blood cell traits are recapitulated (Supplementary Fig. 6a-g). Collectively, SCAVENGE can recapitulate known colocalizations between phenotype-relevant genetic variants and particular cell contexts, while providing additional information enabled by single-cell profiles.

### SCAVENGE uncovers heterogeneity of COVID-19 severity-associated cell populations

A major strength of single-cell approaches is the ability to reveal heterogeneity within an annotated cell type or state, particularly amongst those previously thought to be homogeneous. Given the success of SCAVENGE to identify cell type associations across human hematopoiesis, we were next interested in assessing whether SCAVENGE could capture disease-relevant cell states in phenotypically-rich, but heterogeneous, single-cell data. To this end, we investigated the enrichment of genetic variants associated with increased risk for severe COVID-19 in individuals with a SARS-CoV-2 infection. We employed summary statistics from a GWAS of individuals hospitalized with COVID-19 compared to those who were infected, but not hospitalized, from the COVID-19 Host Genetics Initiative^29^. We performed Bayesian fine-mapping to prioritize risk associations and identified 265 putative causal variants from this COVID-19 severity GWAS^30^ (Supplementary Table 4).

We then applied SCAVENGE to investigate enrichment for these variants using scATAC-seq profiles of 97,315 PBMCs from individuals with moderate or severe COVID-19, as well as healthy donors^31^ (Supplementary Table 5). We observed that the TRS of cells from COVID-19 patients was significantly higher than those from healthy individuals (Supplementary Fig. 7), suggesting that SCAVENGE could capture cell states relevant to disease. We found the monocytes and dendritic cells were enriched for the highest TRS among 15 different cell populations (Fig. 4a,b). This observation is in line with recent reports that different types of monocytes and dendritic cells are intimately associated with inflammatory phenotypes and immune responses in severe COVID-19^32,33,34^.

**Fig.4:**
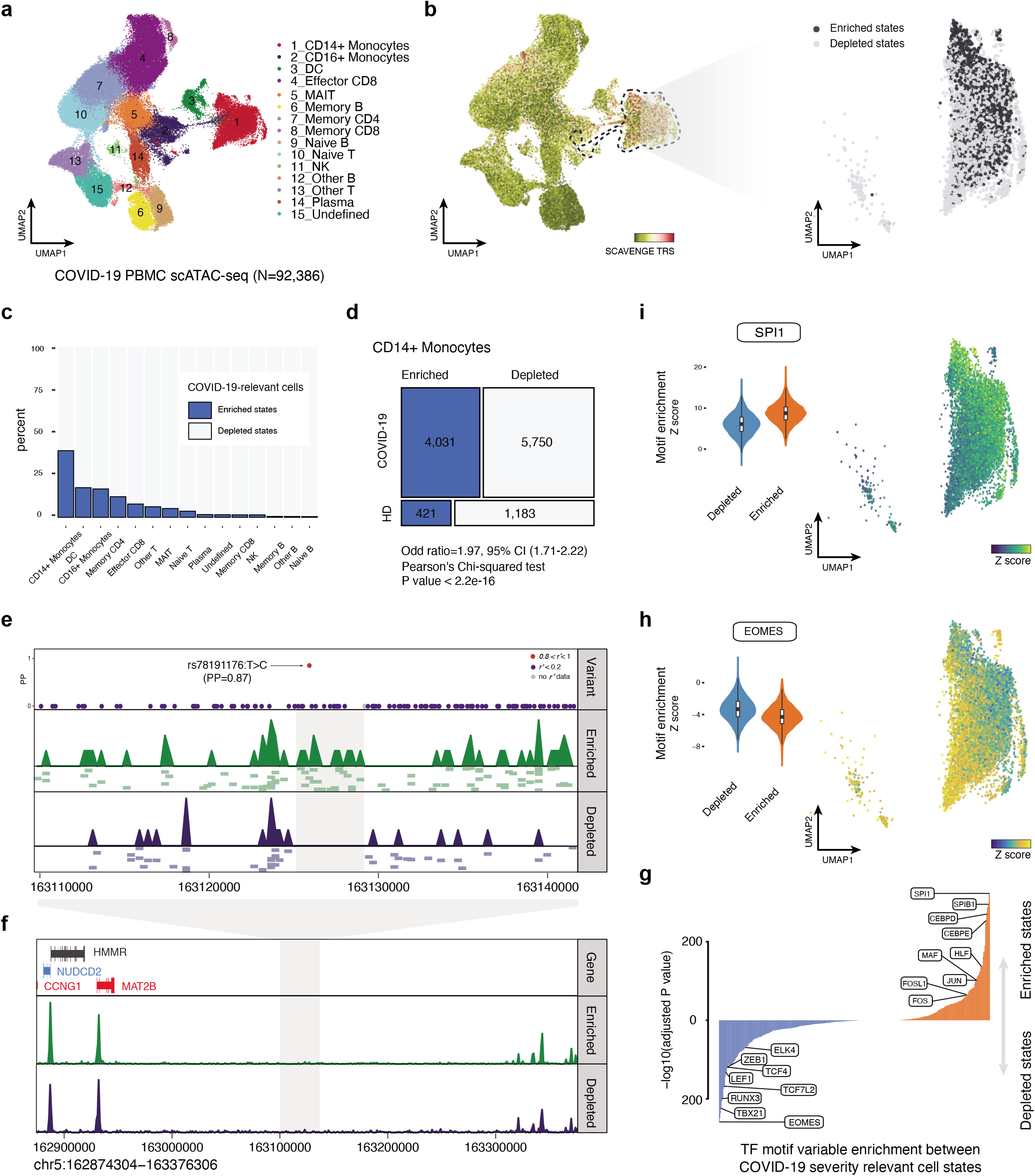
SCAVENGE captures disease-associated cell states and dissects the heterogeneity in association of COVID-19 severity in CD14+ monocytes. **a**, A UMAP plot of scATAC-seq profiles of 92,386 PBMCs from healthy and COVID19 donors^31^. Cells are colored by the cell type annotation. **b**, SCAVENGE TRS for the trait of COVID19 severity risk is displayed for all cells in the UMAP plot (left). CD14+ monocytes are highlighted with dashed lines and two cell states related to COVID-19 risk-variant are identified (Methods) and shown with UMAP coordinates (right). **c**, The percent of cells that are enriched and depleted for COVID19 severity risk variant are shown across all the cell types. **d**, A mosaic plot depicts the distribution of trait-relevant cell states corresponding to disease status. The significance of association is calculated using Pearson’s Chi-squared test. **e-f**, The COVID19 severity variant rs78191176 with high causal probability (PP=0.87) enriched regulatory signals exclusively in trait-relevant cell states (**e**), despite overall a high similarity is observed between pseudo-bulk tracks of these two cell states (**f**). The LocusZoom-style plot of fine-mapped variants is shown and the color represents the degree of linkage (*r*^*2*^). The pseudo-bulk track is aggregated from single-cell accessibility profiles within the same cell state. Further normalization and adjustment is performed to ensure pseudo-bulk tracks can be compared directly. A number of representative single-cell based accessibility profiles are shown below the pseudo-bulk tracks in (**e**). Each pixel represents a 500-bp region. **g**, Differential comparison of chromVAR TF motif enrichment between COVID-19 risk variant-enriched and -depleted CD14+ monocytes. Bonferroni-adjusted significance level is indicated. **h-i**, The chromVAR enrichment Z scores for EOMES (**h**) and SPI1 (**i**) motifs are shown in UMAP plots (right) and violin plots (left) across CD14+ monocytes as shown in (**b**).

We noted marked heterogeneity of risk variant enrichments across different monocyte and dendritic cell populations, which at an aggregated level were the most enriched cell populations for genetic variants conferring risk of severe disease. To dissect this heterogeneity, we performed permutation testing coupled with SCAVENGE to determine an empirical TRS distribution for each cell, instead of using a fixed cutoff (Methods, Supplementary Fig. 8a). This approach enables binary classification of a cell that is enriched or depleted for the trait of interest. Overall, 10.9% of cells were identified as being enriched for variants associated with COVID-19 severity. We focused on the CD14+ monocyte population because this cell type showed the greatest extent of heterogeneity, with 39.1% of cells showing enrichment compared to 5.4% for the other cell types on average (Fig. 4c, Supplementary Fig. 8b). COVID-19 risk variant-enriched cells were significantly associated with diseased cell states in the scATAC-seq data (Fig. 4d, odds ratio (OR) = 1.97, Pearson’s Chi-squared test *P* = 2e-16). By aggregating the cells in identical states into pseudo-bulk populations, we found that the strong association remained in COVID-19 risk-variant enriched cells, but did not hold true for COVID-19 risk-variant depleted cells (enriched state Z = 3.6, depleted state Z = −0.8), suggesting that the risk variants selectively enrich for activity found in individuals with COVID-19 and not in healthy donors. Importantly, we noted key changes in chromatin accessibility in regions where severity risk variants resided, despite the overall profiles being similar (Fig. 4e,f). For instance, a putative causal variant rs78191176:T>C (posterior probability = 0.87) conferring risk for COVID-19 severity overlapped a site showing strong regulatory activity in COVID-19 risk-variant enriched cells, but was absent in COVID-19 risk-variant depleted cells (Fig. 4e). Four genes close to this variant locus were found to be associated with COVID-19. One of these, *MAT2B* was reported as a putative causal gene underlying the genetic risk for severe COVID-19^35,36^, while *CCNG1* was reported as a potential therapeutic target^37^. We found the gene score of both *MAT2B* and *CCNG1* was significantly higher in the enriched cells compared to those from depleted cells (Supplementary Fig. 9), implying that potential regulatory links are mediated by trait-relevant accessible chromatin regions. Critically, our findings point to a likely selective role for COVID-19 severity-associated genetic variants in modulating disease responses in selective cell states, including in immature CD14+ monocyte populations, in the setting of infection with SARS-CoV-2 and that many regulatory elements in which the risk variants reside may not be active in healthy donors.

To define the transcriptional program and transcription factors (TF) that likely drive cell state-specific regulatory programs, we employed chromVAR^38^ to identify TF motif enrichments and examined the differences of single-cell epigenomes between COVID-19 relevant cell states. We observed cell state-specific patterns of TF motif enrichment (Fig. 4g-i). For instance, we observed significantly stronger enrichment of TBX21 (adj *P* =1e10-224), RUNX3 (adj *P* =1e10-199), TCF7L2 (adj *P* =1e10-167), and ZEB1 (adj *P* =1e10-118) motifs in severe COVID-19 risk variant-depleted cells. Many of these factors are critical for regulation of monocyte maturation and serve as markers of non-classical monocytes^39^. In contrast, SP family (SPI1, adj *P* =1e10-320; SPIB1, adj *P* =1e10-295), CEBP family (CEBPD, adj *P* =1e10-270; CEBPE, adj *P* =1e10-248), and AP2 family (JUN, adj *P* =1e10-101; FOS, adj *P* =1e10-51) motifs are prominently more activated in severe COVID-19 risk variant-enriched cells. These TFs play important roles in inflammation-induced myelopoiesis^40^ and monocyte differentiation^41^. Notably, this observation is in line with a recent report^42^ showing that many of these TFs have been implicated in gene expression programs in monocyte subsets associated with COVID-19 disease severity. This observation also implies that these two distinct cell states that are derived from the same cell type (CD14+ monocytes) could be distinctly important to disease. Therefore, SCAVENGE can provide key and previously unappreciated biological insights by taking advantage of the heterogeneity of enrichments for specific disease-associated variants in single-cell data.

### SCAVENGE captures the dynamic risk of predisposition to acute leukemia

While our prior analyses with SCAVENGE have illuminated the ability to identify disease-relevant cell types and states, we wanted to also assess whether disease-relevance across a development trajectory could be achieved. Childhood acute lymphoblastic leukemia (ALL) provides an ideal test case, as the precise cells of origin remain poorly defined in this disease, given that cell state can be altered as a result of malignancy. After fine mapping of the GWAS for risk variants underlying this disease^43,44^ (Supplementary Table 6), the causal variants were employed for SCAVENGE analysis with an scATAC-seq dataset of human hematopoiesis (N=63,882) that we also used in our validation of hematopoietic trait enrichments (Supplementary Fig. 6, Supplementary Table 7). We observed that different types of lymphocytes and their precursors are exclusively enriched for ALL risk associations (Supplementary Fig. 6a, Supplementary Fig. 10). Intriguingly, high enrichments observed in B cell-related populations for ALL are absent in the variety of blood cell traits analyzed, highlighting how genetic effects underlying distinct diseases or traits are mediated through different cellular contexts.

Given existing controversies on the cell of origin for B-cell ALL^45,46^, the most common form of this disease comprising approximately 80% of childhood ALL cases, we focused on assessing the dynamic differentiation process of B cells to explore enrichments across this trajectory. We established a trajectory of B cell development as a continuum from HSCs and progenitors to mature B cells and plasma cells^28,47^ (Fig. 5a). Importantly, the strongest ALL risk variant enrichments were observed across key intermediates in B cell development, including from pro-B cells to naive B cells with a peak in early pre-B cells - a state that is highly relevant to this disease, yet which has not been shown to definitively underlie this disease as a cell of origin (Fig. 5b,c,f). This observed pattern reveals how regulatory chromatin may be impacted by disease-predisposing genetic variants. This analysis also revealed potential mechanisms for specific enriched variants. For instance, a causal variant rs2239630:G>A (posterior probability = 1) located in the core promoter region of *CEBPE*, where regulatory activities undergo dynamic changes across the B-cell developmental trajectory, with the most prominent activity noted at the pre-B cell stage (Fig. 5d). This promoter polymorphism variant was reported as being important for *CEBPE* expression in lymphoblastoid cell lines and the risk allele promotes higher expression of *CEBPE*^*48,49*^. Our findings through SCAVENGE analyses now reveal a specific B cell developmental stage that likely underlies this predisposition.

**Fig.5:**
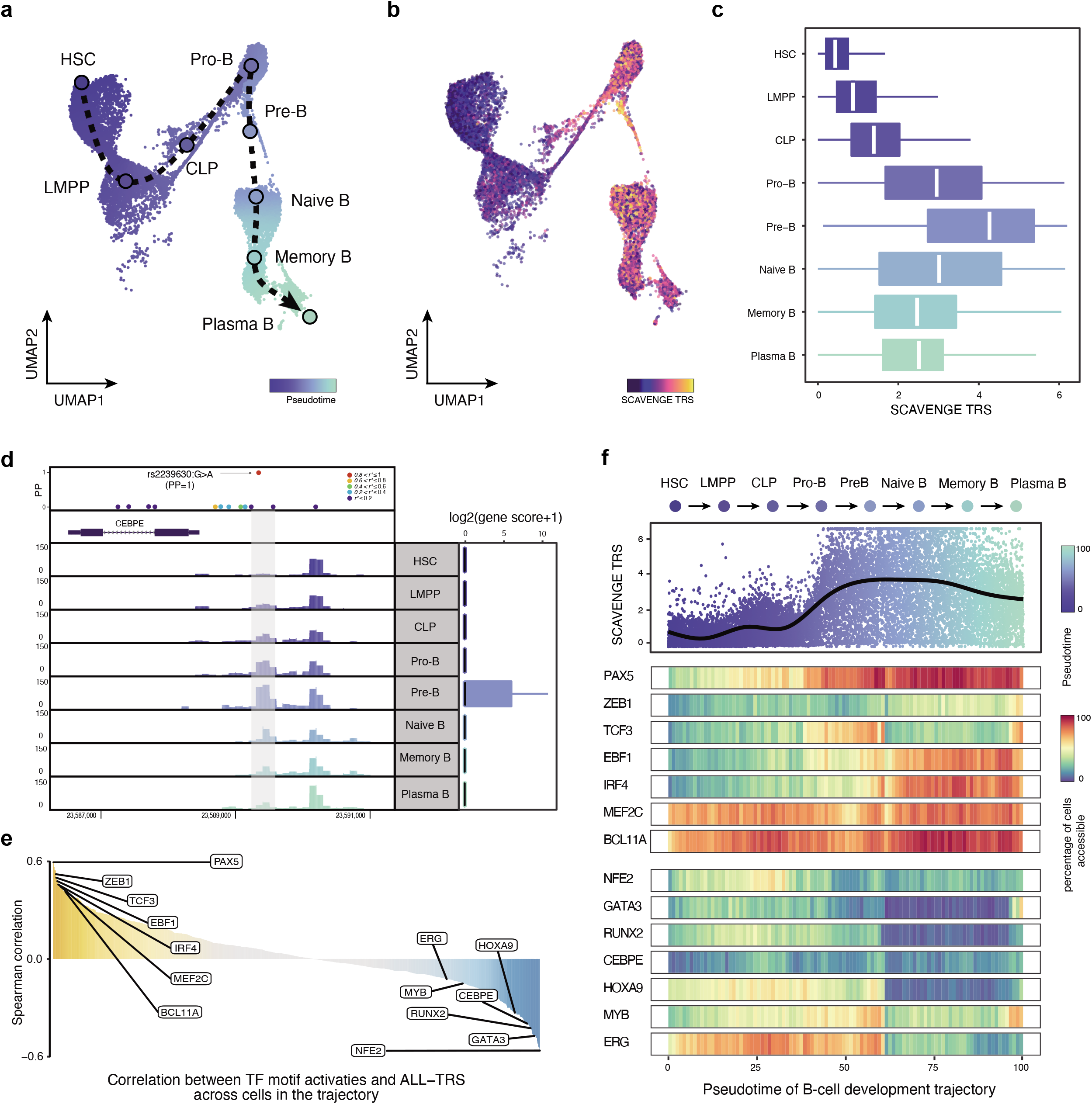
SCAVENGE reveals the dynamic changes of ALL risk predisposition along the B-cell development trajectory. **a**, B-cell differentiation trajectory from HSCs to terminal plasma B cells is constructed. The smoothed arrow represents a visualization of the interpreted trajectory and the cells are colored by pseudo-time. **b-c**, SCAVENGE TRSs of cells are displayed (**b**) in the UMAP embedding and (**c**) box plots across eight development stages in the trajectory. The box plot center line, limits and whiskers represent the median, quartiles and 1.5x interquartile range, respectively. **d**, Regulatory activity is dynamic change along B-cell development stages for the ALL risk variant rs2239630 (PP=0.9996) located in *CEBPE* promoter region, where it reaches peak at pre-B cell stage. Fine-mapped variants are shown in the LocusZoom-style plot and the color represents the degree of linkage (*r*^*2*^). **e**, Spearman correlations between SCAVENGE TRS and chromVAR TF motif enrichment score across all the cells of B-cell development trajectory for TFs from the JASPAR 2018 Database are shown. **f**, SCAVENGE TRSs of cells are depicted along the pseudotime of trajectory (top), the dots are colored by pseudo-time. Pseudotime heatmaps of gene activities for selected TFs that are either positively correlated (middle) or negatively correlated (bottom) are shown. The trajectory is divided into 100 equal bins along the pseudotime. For each bin, we compute the gene activity as the proportion of cells that have non-zero values of gene scores.

To provide further insights and examine which TFs may be relevant to ALL risk across B cell development, we correlated single-cell SCAVENGE TRS with TF motif enrichments across the entire B-cell developmental trajectory. We identified many TFs that are crucial for B cell lineage specification that are positively associated with ALL risk (Fig. 5e,f, Supplementary Fig. 11). Most notably, PAX5 is the most correlated TF, one of the most frequently mutated in ALL^50^ and germline mutations in this TF itself also significantly predispose to ALL acquisition^51,52^, suggesting a unique example where strong predisposition of both variants in a TF and potential *cis*-regulatory targets of the TF may underlie germline cancer predisposition. We also noted a number of TFs that were negatively correlated with risk including CEBPE, GATA3, MYB, and ERG. Notably, several causal variants co-occurred in the vicinity of these TF encoding genes (Supplementary Table 6). These findings suggest that some TFs, such as CEBPE that increases ALL risk with higher expression, may also impact *cis*-acting risk alleles and thereby promote acquisition of ALL. Our findings across the trajectory of normal B cell development provide new insights into the mechanisms underlying predisposition to the most common childhood cancer, B-cell ALL, and suggest opportunities for further developmental and genetic studies of this process using enrichments achieved via SCAVENGE analysis.

## DISCUSSION

Here, we introduce SCAVENGE, a method that characterizes complex disease- and trait-genetic associations in specific cell types, states, and trajectories at single-cell resolution using a network propagation strategy. We demonstrate that SCAVENGE is well-calibrated and powerful through the use of simulated and real datasets. SCAVENGE is robust to parameter choice and reproducible across genetic phenotypes and single-cell datasets. It can be effectively run without requiring parameter tuning or long computation times. Crucially, we have also provided several use cases demonstrating how SCAVENGE can provide previously unappreciated biological and functional insights by mapping disease-relevant genetic variation to an appropriate cellular context.

We anticipate that SCAVENGE will be a valuable tool for discovery of novel cell type/state associations with a range of complex diseases and traits, especially for rare or previously unknown cell populations that are being revealed by the increasing availability of large-scale single cell atlases. While we have focused primarily on use cases involving scATAC-seq datasets here, given their widespread availability, we envision that with the increasing number of other single-cell epigenomic datasets being produced^53,54^ that similar analyses can be conducted with this other data. With the fine-scale causal cell type/state mapping possible with SCAVENGE, the arc of moving from variant to function can start to be filled in a systematic and precisely targeted manner. For instance, functional experiments can be directly performed in the most relevant cell populations. In combination with additional inference tools, including the use of computation approaches to predict target genes of particular *cis*-regulatory elements^13,55,56^, as well as other functional genomic data, including chromatin conformation (e.g. Hi-C)^57^ and TF occupancy data (e.g. CUT&Tag)^58^, prediction of how particular variants could alter these regulatory elements, modify gene expression, and result in human disease will become possible - the ultimate goal of moving from variant to function.

SCAVENGE offers a versatile framework that can be easily adapted to other single-cell genomic analyses, given that high sparsity remains a central challenge in diverse single-cell modalities (e.g. 55–90% of expressed genes suffer from dropout in single-cell RNA-seq data sets^15^). As single-cell genomic datasets grow in volume, SCAVENGE holds great promise for efficiently uncovering relevant cell populations for more phenotypes or functions in different scenarios, which may expand beyond the complex trait genetic variants we have examined here. We envision the SCAVENGE framework can enable critical new biological insights, akin to the way that search engines such as Google have accelerated our ability to find relevant information across the vast sea of information on the internet.

## Methods

### SCAVENGE methodology

The framework of the SCAVENGE method is schematized in Fig. 1. Briefly, SCAVENGE employs a network propagation strategy to explore transitive associations of a subset of cells that are highly relevant to traits of interest in the cell-to-cell network. The workflow is described in detail in the following steps:

#### Cell-to-cell similarity network construction

SCAVENGE uses a mutual *k* nearest neighbor (M-*k*NN) graph to faithfully represent inherent relationships of individual cells. We start from the feature-by-cell matrix from scATAC-seq profiles and use latent semantic indexing (LSI)^59,60,61^ to extract representative lower-dimensions. Specifically, the binarized sparse matrix is first converted into a term frequency-inverse document frequency (TF-IDF) matrix by weighting the matrix against the total number of features for each cell with the following formula:

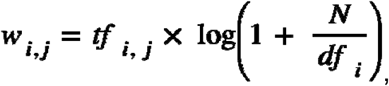

where ***w***_***i,j***_ is the weight for the feature ***i*** in cell ***j***, ***tf***_*i,j*_, indicates the term frequency that is the number of feature ***i*** in cell ***j***, ***df***_***i***_ is the document frequency of term that is number of cells where the feature ***i*** appears, ***N*** is the total number of cells in the experiment.

The singular value decomposition (SVD) is applied on the TF-IDF matrix to generate an LSI score matrix with a lower-dimensional space as follows:

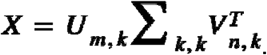

This is the decomposition of ***X*** where ***U*** and ***V*** are orthogonal matrices and ***Σ*** is a diagonal matrix. ***m*** represents the number of rows and ***n*** represents the number of columns for ***X***. ***U*** = [***μ***_**1**_, … , ***μ***_***k***_] is the left singular vector and ***μ***_1_ with length ***m***. ***V*** = [***ν***_**1**_, … , ***ν***_***k***_] is the right singular vector and ***ν***_***i***_ with length ***n***. ***Σ***_***k, k***_ = ***diag***(***σ***_**1**_, … , ***σ***_***k***_) and ***σ***_**1**_ ≥ ***σ***_**2**_ ≥ … ≥ ***σ***_***k***_ are singular values of ***X***.

Next, SCAVENGE builds a nearest neighbor graph from the LSI matrice of cells and leading LSIs (***d***=30). The Euclidean distance between any pair of cells is calculated based on the LSI matrix and *k* nearest neighbors (*k*=30) for each cell are identified. To rigorously ensure cells connected with the same phenotype or state, we construct an M-*k*NN graph by requiring the node (cell) pair in the graph that are mutually the *k* nearest neighbors to each other^18,62^. If a cell fails to find its mutually *k* nearest neighbors, we connect this cell to its nearest neighbor to guarantee the connectivity of the resulting graph. The M-*k*NN graph enables prioritization of *k*NN structure and allows each cell to have at most *k* neighbors. It could efficiently avoid the generation of extreme hubs that have a large number of neighbors and also ensure the sparsity of the resulting graph^18,62^.

#### Definition of seed cells

Given the sparsity of single-cell genomic data, only a few of cells in the top ranks evaluated by the colocalization approaches could reliably reflect the relevance to a phenotype/trait/disease of interest (Supplementary 1a,b, Fig. 2a). We define the seed cells as a set of cells that are most likely to be relevant to the tested trait. To identify the seed cells for a specific trait of interest, we use g-chromVAR to calculate a bias-corrected Z score (confounding technical factors such as GC content bias and PCR amplification) for each cell^13^, by integrating posterior probabilities of genetic causal variants and their strength of chromatin accessibility. We realized that seed cells and their number can vary among tested genetic traits and the single-cell dataset employed, it is not suitable to pre-define a fixed number for seed cells that is optimal for all situations. As the Z score generated from g-chromVAR is a normalized measurement that considers cells uniformly across all the cells, we reasoned that this can serve as an initial filter for seed cells. We convert Z scores to *P* values using a one-tailed normal distribution and initially consider all the cells with P value less than 0.05 as seed cells. Our analysis showed that SCAVENGE is robust to a range of proportions for seed cell selection (Supplementary Fig. 4a). In practice, 5% of total cells is a number sufficient to represent seed cells. We therefore refine the seed cells by keeping the 5% cells with the highest Z score if the number of initial seed cells exceeds 5%.

#### Network propagation with seed cells

SCAVENGE relies on the concept of network propagation, which is based on the Guilt By Association principle where the proximity between the set of seed nodes and all the nodes in the graph can be comprehensively measured. We introduce random walk with restart (RWR)^63^, a network propagation-based algorithm for propagating the set of seed cells for a trait of interest to efficiently discover the transitive associations hidden in the cell-to-cell graph.

In general, a random walk over a graph is a stochastic process. That means the initial state of the graph is known and its state changes over iterations (random walk) with a transition probability matrix that describes the probability for one node jumping to another. The initial state of the graph is defined by selected seed nodes (cells). Their information propagates to all nodes in the graph and the graph finally reaches a stationary state after a series of random walk processes. The strength of transitive associations can be measured by information carried by each node at the stationary state of the graph. The stationary distribution is defined as the network propagation score. By leveraging the entire structure of the graph, the RWR algorithm allows the measurement of a cell influenced by seed cells from not only its direct neighborhood but also the distant immediate neighborhood that can be reached by multiple steps. Intuitively, the more a cell is influenced by the seed cells, the greater relevance it has to the phenotype/trait evaluated. As such, a higher network propagation score indicates stronger relevance to the trait evaluated.

More formally, there is a set of seed nodes ***I* ∈ *V*** defining the initial states of an undirected M-*k*NN graph, ***G* = (*V*, *E*)** that is constructed as described above. Two sparse matrices are created to represent graph structure.

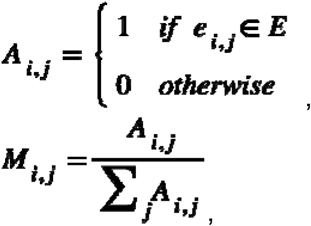

where ***A***_***i,j***_ denotes adjacency matrix of ***G***, ***M***_***i,j***_ is transition probability matrix that is column normalization of ***A***_***i,j***_.

The random walk steps ***s*** is discrete and finite, ***s* ∈ N**. The information carried of node ***v*** at step ***s*** is ***v***_***s***_. We considered that the information is equally distributed across the seed nodes in the initial state of step 0. We can write

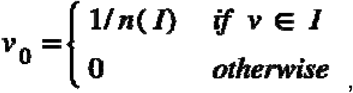

where ***n*(*I*)** is the number of seed nodes.

At each iteration, a node can transfer the information to one of its randomly selected neighbors (the probability is proportional to the number of neighbors and stored in ***M***_***i,j***_) or restart at the node by transferring information back to itself. As the random walk is irreducible and aperiodic, the iterative update of this procedure is guaranteed to converge to the stationary steady-state. The corresponding stationary distribution or probability for each node can be obtained by recursively applying the following equation until convergence: ***∀ i***, ***j ∈ V***, ***∀ s ∈* N**,

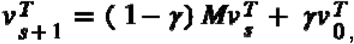

where ***γ*** is the restart probability ranged from 0 to 1 (***γ***=0.05). A restart probability in practice can avoid the walk being trapped in a dead-end, and assure the quick convergence of the graph to achieve the stationary distribution. The random walk process is continued until steady-state (|***ν*** − ***ν***_***s***_**| < *α***, where ***α*** is 1e-5) and stationary distribution ***ν***_***s***_ is considered as the network propagation score.

#### TRS normalization and scale

The total sum of information is kept constant and equals 1 throughout the graph while information spreads over the graph with each iteration. As such, the network propagation score averaged per cell is proportional to the cell number we evaluated. Therefore the original network propagation scores are essentially very small and can not be compared with different datasets, even if the same genetic trait is assessed. At the same time, we cannot determine significance for each cell from the network propagation score. We reasoned that appropriate processing of network propagation scores is needed to enable per-cell scores to be comparable across different traits, but inherit the overall significant levels from our g-chromVAR analysis. To this end, we define the trait relevance score (TRS) by scaling and normalizing the network propagation score. Specifically, we first calculate ***gg***^***th***^ percentile of network propagation scores and use it as the ceiling. This step makes sure that a few cells with high network propagation scores are put on the same level and avoids the impacts of potential extreme outliers so that network propagation scores could be efficiently scaled up between 0 and 1 across all cells.

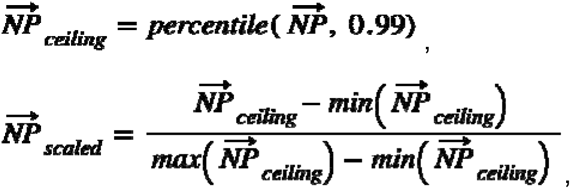

To match the final TRS to the significant level of the original dataset, a scaling factor representing the average levels of bias-corrected Z score with the top 1% of cells is calculated by the following:

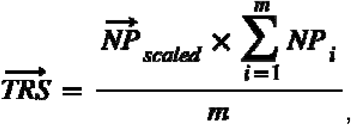

where m is a cell with the top 1% bias-corrected Z score.

#### Cell state identification using the permutation test

To further assess whether a cell is enriched or depleted for the trait of interest, we propose a method to determine the statistical significance for individual cells by calculating an empirical distribution of scores per cell, instead of using a fixed cutoff arbitrarily (Supplementary Fig. 8a). Here we focus on network propagation score rather than TRS because network propagation scores directly result from network propagation processes without further normalization and scale, the sum of network propagation scores across all cells is constantly equal to 1 and independent of seed cell selection, such that the network propagation scores yielded from SCAVENGE analysis with different sets of seed cells are directly comparable. We use a permutation-based method to generate the empirical distribution. We randomly select a set of number-matched seed cells to repeat SCAVENGE analyses. To maintain consistency of topology attributes with real seed cells, we require the permuted seed cells to have the same degree distribution for each permutation. That means if a seed cell has *m* neighbors in the cell-to-cell graph, the matched permuted one can only be selected from cells with *m* neighbors. The enriched cell is expected to have a larger network propagation score than that from permuted seed cells. The permutations can be repeated independently multiple times. For each cell, the significance can be determined by comparisons of real network propagation scores and those from permutations. The empirical *P* value is defined as ***∀***_***c***_   {***cell***_**1**_, **… *cell***_***N***_**}**,

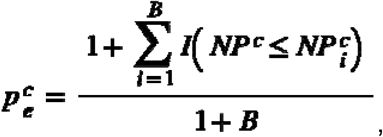

where ***B*** is the number of permutations (***B***=1,000). The empirical *P* value is calculated as the proportion of the network propagation score of permutation greater than its real score. Trait-enriched cells are defined as cells with *P* less than 0.05

### Assessing performance with simulations

To evaluate the calibration and power of our method, we conducted simulations from downsampling a variety of FACS-sorted bulk hematopoietic populations^13^. The simulation framework utilizes an approach that has been described previously^15^. We started from a peak-by-cell count matrix generated from bulk ATAC-seq data. The read count of synthetic single cells for peak *i* in cell type *t* follows a binomial distribution 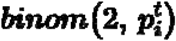, where 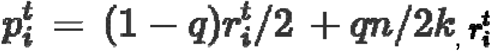 is the ratio of reads for peak *i* in cell type *t* from the bulk ATAC-seq data; *k* is the total number of peaks in the bulk data and *n* is the number of simulated fragments; *q* specifies noise level (***q* ∈ [0. 1]**), where *q*=0 is no noise and *q*=1 indicates the highest level of noise, which means a random distribution of n fragments into *k* peaks.

We used g-chromVAR to assess enrichment of the highly heritable trait of monocyte count across 16 hematopoietic cell types (Supplementary Fig 3). For ease of interpretation and validation, we created a ground truth dataset including monocyte and NK cells to represent enriched and depleted cell populations, respectively. We simulated 500 cells per labeled cell type with the parameters of *n*=10,000 and *q*=0.3. Eventually, 1,000 cells were simulated. Intuitively, the top ranked 500 cells will be monocytes while bottom 500 ranked cells will be NK cells if these cells are perfectly classified. Additional simulations are also generated to investigate the robustness of SCAVENGE. We set parameter *n* to various values including 2,500, 5,000, 7,500, 10,000, 25,000 and 50,000 to test the effects of sequencing depth. We set *q* to various values including 0.25, 0.3, 0.35 and 0.4 to test the robustness to noise. To qualitatively assess the performance of SCAVENGE in a more complex situation, we also generated another dataset consisting of 9 cell types that showed trait relevance at different levels, where 200 cells per labeled cell type were synthesized. SCAVENGE was applied to these simulated datasets using the default parameters, except for evaluation of the number of seed cells and number of neighbors used for graph construction.

### Application of SCAVENGE to the scATAC-seq datasets

Four independent datasets were used for SCAVENGE analysis as use case examples in this study. All cell-type annotations and metadata were obtained from the original studies unless we specifically state otherwise below.

#### The 10X Genomics Peripheral Blood Mononuclear Cell (PBMC) dataset

We downloaded fragment files of this dataset from the 10X Genomics website (https://support.10xgenomics.com/single-cell-atac/datasets/1.0.1/atac_v1_pbmc_5k). This PBMC dataset includes 5,335 cells from one donor and no cell annotations are provided. The dataset was processed by the standard ArchR pipeline with default parameters^60^, including Arrow files creation, quality control, inferring doublets, dimensionality reduction, and clustering. We initially obtained 8 cell clusters and kept 6 for those containing at least 50 cells in each cluster. We retained 4,562 cells for SCAVENGE analysis for the trait of monocyte count. Gene scores of several cell type-specific marker genes were calculated based on chromatin accessibility in the vicinity of the gene and used to annotate PBMC populations.

#### Hematopoiesis ATAC-seq dataset

This hematopoietic cell dataset consists of 35,038 cells from 2 bone marrow mononuclear cell (BMMC) donors, 3 CD34+ enriched bone Marrow Cells (CD34+) donors, and 5 PBMC donors. The data were processed as described in the original publication^23^. We downloaded the processed data with summarized experiments format from https://github.com/GreenleafLab/MPAL-Single-Cell-2019. Three cell types were removed owing to unknown cell labels. A total of 33,819 cells from 23 cell populations were selected for further analysis. The LSI-by-cell matrix with the first 30 leading LSIs is extracted for M-*k*NN graph construction. The peak-by-cell matrix is used as input for SCAVENGE analysis for 22 blood cell traits. The per-cell-based TRS is visualized with uniform manifold approximation and projection (UMAP)^64^ coordinates. For each tested blood trait, the TRSs from the same cell population collapsed into the median value to represent the TRS on the cell-type level.

#### Hematopoiesis scATAC-seq dataset 2

This hematopoietic dataset consists of 63,882 cells from 1 BMMC donor, 2 CD34+ donors, and 16 PBMC donors. The data were processed as described in the original publication^28^. We downloaded the processed peak-by-cell count matrix as well as cell annotations of this dataset from https://github.com/GreenleafLab/10x-scATAC-2019. A total of 63,882 cells of 31 cell types were used for SCAVENGE analysis for a variety of blood cell traits, which is similar to the above hematopoiesis scATAC-seq dataset. We also applied SCAVENGE on this dataset to explore the enrichment of ALL associations.

We also construct a single-cell trajectory of B-cell development to further examine how ALL risk is variably enriched along this trajectory. This trajectory consists of 8 cell types from HSCs to progenitors to mature B cells. The pseudotime for each cell in this trajectory was calculated as previously described^28^. To identify transcription factors correlated with genetic trait enrichments, we calculated the Spearman correlation between TF motif enrichment scores and SCAVENGE TRSs using all cells in the trajectory. The trajectory was divided into 100 equal bins along the pseudotime. For each bin, we computed the gene activity as the proportion of cells that have non-zero values of gene scores. Gene activities for selected TFs were shown in the pseudotime heatmaps.

#### COVID-19 PBMC scATAC-seq dataset

This dataset comprises 97,315 PBMCs, obtained from 3 healthy donors and 8 COVID-19 patients, of which 5 had moderate disease and 3 had severe disease. The fragment files processed using the Cell Ranger pipeline were obtained from the authors of the original paper^31^. We performed cell clustering and cell-type annotation using the ArchR package^60^. We created arrow files from fragment files and performed quality control with metrics including the number of unique fragments and enrichment of the transcription-start-site. Iterative latent semantic indexing was performed with ‘addIterativeLSI’ function and potential sample-specific and batch effects are corrected using the harmony algorithm with ‘addHarmony’ function. We applied UMAP dimensionality reduction and Leiden clustering^65^ to the batch-corrected epigenomic datasets. Initially, 25 cell clusters were identified and we merged similar cell clusters and annotated cell populations using gene scores of canonical markers from the original publication. For the resulting 15 cell types, we performed peak calling and generated the peak-by-cell matrix for SCAVENGE analysis of the COVID-19 severity associated genetic variants.

To explore the heterogeneity of trait-associated enrichments, we performed cell state discovery analyses as described above and the cells were segregated into a severe COVID-19 risk variant-enriched population and a severe COVID-19 risk variant-depleted population. The number and proportion of these two cell states were investigated across individual cell types. We found that in most cell types, the cell number across these two cell states are extremely different. We thus selected the same amount of cells that are most representative of each cell state for further analysis. In the case of CD14+ monocytes, 1,000 cells with the highest TRS in severe COVID-19 risk variant-enriched cells and 1,000 cells with the lowest TRS in severe COVID-19 risk variant-depleted cells are selected to explore the differences of TF motif enrichment in the peak region. The accessibility profiles of these 2,000 cells were used to compute gene scores for genes of interest. The corresponding genome browser accessibility tracks of single-cell-based occupancy and pseudo-bulk samples are plotted using the ‘plotBrowserTrack’ function.

### GWAS summary statistics and fine-mapping analysis

#### Blood cell traits

Summary statistics of 22 blood cell traits from Blood Cell Consortium 2 (BCX2) were processed as previously described^22^. Variants with fine-mapped posterior probability > 0.001 for a locus in one or more blood traits were retained and used for analysis.

#### COVID-19 severity

We obtained summary-level GWAS data of B1 (hospitalized COVID-19+ versus non-hospitalized COVID-19+) from the COVID-19 Host Genetics Initiative (release 5, https://www.covid19hg.org) with restricted ancestry to EUR individuals. This COVID-19 severity trait is from a meta-analysis of 13,641 moderate or severe COVID-19 hospitalized cases and 49,562 reported cases of SARS-CoV-2 infection. Given that only summary level data were available in this instance (raw genotype level data were not accessible), conditionally independent signals were first identified using GCTA-COJO^66^. In COJO, window size was set to 10Mb, p-value threshold was set to a suggestive level of 1.0e-06 because of limited signal reaching genome-wide significance. Subsequently, approximate Bayesian factor (ABF) analyses were performed as described^30^ using a window size of 1Mbp on either side of independent variants. The prior variance in allelic effects was estimated as 0.04, considered to be broadly appropriate for this method, and calculated using formula (8)^30^. For loci containing multiple independent signals, association statistics surrounding an index variant in question were based on corrected GCTA approximate conditional analysis adjusting for all other independent variants in that 1Mbp either side region. Finally, the posterior probability of being causal (PP) was calculated by dividing the ABF of each variant by the sum of ABF values over all variants in the window. LocusZoom-style plots were created in R, using a 1000G EUR-subsetted reference panel for LD information.

#### Acute Lymphoblastic Leukemia predisposition

The GWAS data of childhood ALL was obtained from our previous study^44^. For causal variant identification, we performed fine-mapping at 13 well-replicated and 3 novel ALL risk loci identified in our recent trans-ancestry GWAS. In this instance, where raw genotype data were available, FINEMAP was used^67^. An LD matrix was created for 1Mbp either side of lead significant variants using an unrelated set of genotypes (3rd-degree-relatives or closer), including all ancestry groups. FINEMAP was run in the stochastic search method, with all defaults in place apart from --n-causal-snps=10 and the posterior probabilities of variants being causal were obtained. Due to significant overlap at the *BMI1*-*PIP42A* locus, variants contributing more causal information (higher PP) were preferentially included.

### The sparsity of scATAC-seq

To assess the sparsity of scATAC-seq data, we used 5 published datasets including 10X PBMCs (N=4,562), leukemic cells (Leukemia, N=391)^68^, a mixture of GM12878 and HEK293T cells (GM12878vsHEK, N=526)^59,68^, a mixture of GM12878 and HL-60 cells (GM12878vsHL, N=597)^59^, and a mixture of breast tumor 4T1 cells (Breast_Tumor, N=384)^69^. These datasets cover two commonly used scATAC-seq platforms of microfluidics (10X PBMCs, Leukemia, Breast_Tumor) and cellular indexing (GM12878vsHEK, GM12878vsHL). The 10X PBMCs dataset was obtained and processed as described above. The other four datasets were processed as previously reported^70^. We downloaded the h5ad files from https://github.com/jsxlei/SCALE, and extracted peak-by-cell matrices, respectively. Two measures of sparsity were examined, 1) the sparsity of peaks, which indicates what proportion of cells have an absence of signal for a given peak; 2) the sparsity of cells, which indicates what proportion of peaks have an absence of signal for a given single cell (Supplementary Fig 1). Because the peak calling is performed with a pseudo-bulk sample, which is generated by the aggregation of all single-cell profiles in each dataset, which implies every peak will present abundant signals in pseudo-bulk data. As the pseudo-bulk accessibility data is highly correlated to and resembles a bulk ATAC-seq experiment, we thus reasoned that these two measurements could well represent the sparsity of individual cells compared to corresponding bulk or pseudo-bulk ATAC-seq data.

### Transcription factor motif analysis

We used chromVAR^38^ to measure global TF activity. We used peak-by-cell matrix and transcription factor motifs within the non-redundant JASPAR 2018 CORE vertebrate (*N*=322) to compute bias-corrected deviation Z score for each cell. We compared motif enrichment Z scores of cells with variable states by using Benjamini–Hochberg-corrected *P* values from one-sided Student’s t-test.

## Data availability

All single-cell datasets used in the paper are public. The processed and analyzed data are publicly available at (https://github.com/sankaranlab/SCAVENGE-reproducibility). Blood cell trait summary statistics are available from (http://www.mhi-humangenetics.org/en/resources/). COVID-19 severity summary statistics are available at (https://www.covid19hg.org/).

## Code availability

SCAVENGE is implemented as an R package and is available on Github (https://github.com/sankaranlab/SCAVENGE). The code to reproduce the results is available on a dedicated GitHub repository (https://github.com/sankaranlab/SCAVENGE-reproducibility).

## Acknowledgments

We are grateful to members of the Sankaran laboratory and numerous colleagues for valuable comments and suggestions. We are thankful to Pengyuan Yang and Maojun You for sharing processed fragment files of PBMC scATAC-seq data. This work was supported by the New York Stem Cell Foundation (V.G.S.), a gift from the Lodish Family to Boston Children’s Hospital (V.G.S.), and National Institutes of Health Grants R01 DK103794 and R01 HL146500 (V.G.S.).

J.S.W. is an Investigator of the Howard Hughes Medical Institute. V.G.S. is a New York Stem Cell-Robertson Investigator.

## Author contributions

F.Y. and V.G.S. designed and conceptualized the study. F.Y. designed and implemented the algorithm and analyzed the data. L.D.C., C.W., L.A.L., S.J., K.X., C.W.K.C., J.L.W., J.S.W., and A.J.d.S., processed the GWAS and fine-mapping data and interpreted the results. F.Y. and V.G.S. wrote the manuscript with input from all authors. V.G.S. provided project oversight.

## Corresponding author

Correspondence and requests for materials should be addressed to Vijay Sankaran.

## SUPPLEMENTARY FIGURE LEGENDS

**Supplementary Fig. 1: The extensively high sparsity in scATAC-seq data**. The density plots show that high sparsity commonly exists across curated scATAC-seq data. The sparsity in scATAC-seq data is characterized using the peak-by-cell matrix from five different datasets. **a**, The sparsity of peaks is defined as the proportion of cells that show no signal (zero-valued) for a given peak. **b**, The sparsity of cells is defined as the proportion of peaks that show no signal for a given cell. The 10X PBMC scATAC-seq dataset is used in the following SCAVENGE analysis.

**Supplementary Fig. 2: Challenges for identification of trait/phenotype-relevant cells using colocalization-based approaches and the network-based solution**. To investigate the causal cell type/state that is relevant to a genetic trait, the most commonly used strategy is co-localization of epigenetic signals that occur in regulatory elements (peaks) and risk variants. However, this approach is unfortunately uninformative for a majority of cells when applied to scATAC-seq profiles. Given the noise and sparse nature of scATAC-seq data, absence of signals are extensive across cells and regulatory peaks, which can not be distinguished between technical or biological causes. Therefore, only a few cells demonstrate reliable phenotypic relevance (**a**). While global high-dimensional features of individual single cells are sufficient to represent the underlying cell identities or states, which enables the relationships among such cells to be readily inferred. We reason that the real relevant cell populations can be revealed and recovered by building a search engine for cell-to-cell networks that enable discovery of similar cells with the same phenotype (**b**).

**Supplementary Fig. 3: Enrichments of monocyte count associated genetic variants in bulk hematopoietic ATAC-seq data**. The enrichment scores are obtained by using g-chromVAR for the trait of monocyte count with bulk ATAC-seq profiles across 16 hematopoietic cell types.

**Supplementary Fig. 4: SCAVENGE is robust to a range of parameter choices and data sparsity and noise. a-b**, The same trait and simulated dataset used in **Fig. 2a** are employed to explore the effects of SCAVENGE performance from (**a**) different fractions of cells selected as seed cells and (**b**) different numbers of *k* used for cell-to-cell network construction. **a**, True positive rate (TPR) is calculated for SCAVENGE analysis by selecting different fractions from 1% to 30% of top ranked cells as seed cells. The red dash line indicates the TPR without SCAVENGE. **c-d**, Different simulated datasets are created to test if SCAVENGE is robust to data sparsity and noise. **c**, The bar plots depict TPRs for simulated data created from different numbers of fragments subsampled from the bulk-level data with (dark color) and without (light color) SCAVENGE. **d**, The bar plots depict TPRs for simulated data created with different levels of noise with (dark color) and without (light color) SCAVENGE.

**Supplementary Fig. 5: SCAVENGE identifies trait-relevant cell populations on a real scATAC-seq dataset. a**, UMAP plots display M-*k*NN graph construction from the latent space for the 10X PBMC scATAC-seq profiles. **b**, UMAP plots illustrate the canonical marker genes of monocytes (*S100A12*, *MPO*), B cells (*PAX5*, *MS4A1*), and T cells (*CD7*, *CD3D*). The cells are colored by corresponding gene activity score.

**Supplementary Fig. 6: SCAVENGE analysis of 22 blood cell traits in a hematopoiesis scATAC-seq dataset 2**. The figures here are similar to those displayed in **Fig. 3**. **a**, UMAP demonstrates the cell type labels. **b, d-g**, Per-cell TRS for four representative traits are shown in UMAP plots (**b**) and per cell type (**d-g**). Boxplots show the median with interquartile range (IQR) (25–75%); whiskers extend 1.5x the IQR. **c**, The median TRSs of cells belonging to the same cell type are shown in the heatmap. Unsupervised clustering analysis is performed and traits are grouped into four clusters using the *K*-means clustering algorithm.

**Supplementary Fig. 7: SCAVENGE captures cell states relevant to COVID-19**. Box plots depicting SCAVENGE TRS of COVID19-severity risk for cells from healthy individuals and patients with mild and severe COVID-19. The significance was calculated using the Student’s t-test.

**Supplementary Fig. 8: Identification of cell states relevant to COVID-19-severity risk. a**, An illustration of the method used to calculate empirical *P* values for cell state determination. The background network propagation score is calculated based on SCAVENGE analysis using randomly selected seed cells that match topology attributes of real seed cells. A *null* distribution of network propagation score is generated and the empirical *P* value is calculated from comparison between network propagation score and *null* distribution (Methods). **b**, Cell number distribution of COVID-19 severity risk variant-enriched and -depleted cell states across all the cell types. Barplots depicting cell numbers and the order of cell types are kept identical to that from **Fig. 4c**.

**Supplementary Fig. 9: Chromatin activity scores of genes in the vicinity of cell state-associated risk variants**. Log-normalized gene scores between COVID-19 severity risk variant-enriched and -depleted cell populations for indicated genes are shown.

**Supplementary Fig. 10: SCAVENGE analysis of ALL risk predisposition**. Per cell SCAVENGE TRS of ALL risk predispositions are demonstrated in the UMAP embeddings of Hematopoiesis scATAC-seq dataset 2. Cell type labels are shown in **Supplementary Fig. 6a**.

**Supplementary Fig. 11: Correlations of SCAVENGE TRS and TF motif enrichment score**. Scatter plots show TFs including (**a**) PAX5 and (**b**) EBF1 that are positively correlated with ALL risk predisposition and TFs including (**c**) NFE2 and (**d**) RUNX2 that are positively correlated with ALL risk predisposition. The dots are colored by TRS. Spearman correlation is calculated between SCAVENGE TRS and chromVAR TF motif enrichment score across all the cells of B-cell development trajectory.

## SUPPLEMENTARY TABLES

**Supplementary Table 1: SCAVENGE analysis of monocyte count with 10X PBMC scATAC-seq dataset.**

**Supplementary Table 2: SCAVENGE analysis of 22 blood cell traits with Hematopoiesis scATAC-seq dataset.**

**Supplementary Table 3: SCAVENGE analysis of 22 blood cell traits with Hematopoiesis scATAC-seq 2 dataset.**

**Supplementary Table 4: Fine-mapped variants of COVID19 severity trait.**

**Supplementary Table 5: SCAVENGE analysis of COVID19 severity with COVID-19 PBMC scATAC-seq dataset.**

**Supplementary Table 6: Fine-mapped variants of the ALL risk trait.**

**Supplementary Table 7: SCAVENGE analysis of ALL risk predispositions with Hematopoiesis scATAC-seq 2 dataset.**

## REFERENCES

1. Cano-Gamez, E. & Trynka, G. From GWAS to Function: Using Functional Genomics to Identify the Mechanisms Underlying Complex Diseases. Front. Genet. 11, 424 (2020).

2. Schaid, D. J., Chen, W. & Larson, N. B. From genome-wide associations to candidate causal variants by statistical fine-mapping. Nat. Rev. Genet. 19, 491–504 (2018).

3. Finucane, H. K. et al. Partitioning heritability by functional annotation using genome-wide association summary statistics. Nat. Genet. 47, 1228–1235 (2015).

4. Rozenblatt-Rosen, O. et al. Building a high-quality Human Cell Atlas. Nat. Biotechnol. 39, 149–153 (2021).

5. Regev, A. et al. The Human Cell Atlas. Elife 6, (2017).

6. Website. https://commonfund.nih.gov/HuBMAP.

7. Domcke, S. et al. A human cell atlas of fetal chromatin accessibility. Science 370, (2020).

8. Zhang, K. et al. A single-cell atlas of chromatin accessibility in the human genome. Cell 184, 5985–6001.e19 (2021).

9. Soskic, B. et al. Chromatin activity at GWAS loci identifies T cell states driving complex immune diseases. Nat. Genet. 51, 1486–1493 (2019).

10. Mahajan, A. et al. Fine-mapping type 2 diabetes loci to single-variant resolution using high-density imputation and islet-specific epigenome maps. Nat. Genet. 50, 1505–1513 (2018).

11. Farh, K. K.-H. et al. Genetic and epigenetic fine mapping of causal autoimmune disease variants. Nature 518, 337–343 (2015).

12. Chiou, J. et al. Single-cell chromatin accessibility identifies pancreatic islet cell type- and state-specific regulatory programs of diabetes risk. Nat. Genet. 53, 455–466 (2021).

13. Ulirsch, J. C. et al. Interrogation of human hematopoiesis at single-cell and single-variant resolution. Nat. Genet. 51, 683–693 (2019).

14. Calderon, D. et al. Landscape of stimulation-responsive chromatin across diverse human immune cells. Nat. Genet. 51, 1494–1505 (2019).

15. Chen, H. et al. Assessment of computational methods for the analysis of single-cell ATAC-seq data. Genome Biol. 20, 241 (2019).

16. Brin, S. & Page, L. The anatomy of a large-scale hypertextual Web search engine. Computer Networks and ISDN Systems vol. 30 107–117 (1998).

17. Cowen, L., Ideker, T., Raphael, B. J. & Sharan, R. Network propagation: a universal amplifier of genetic associations. Nat. Rev. Genet. 18, 551–562 (2017).

18. Levine, J. H. et al. Data-Driven Phenotypic Dissection of AML Reveals Progenitor-like Cells that Correlate with Prognosis. Cell 162, 184–197 (2015).

19. Setty, M. et al. Characterization of cell fate probabilities in single-cell data with Palantir. Nat. Biotechnol. 37, 451–460 (2019).

20. Moon, K. R. et al. Visualizing structure and transitions in high-dimensional biological data. Nat. Biotechnol. 37, 1482–1492 (2019).

21. Aldous, D. J. Lower Bounds for Covering Times for Reversible Markov Chains and Random Walks on Graphs. (1988).

22. Vuckovic, D. et al. The Polygenic and Monogenic Basis of Blood Traits and Diseases. Cell 182, 1214–1231.e11 (2020).

23. Granja, J. M. et al. Single-cell multiomic analysis identifies regulatory programs in mixed-phenotype acute leukemia. Nat. Biotechnol. 37, 1458–1465 (2019).

24. Liggett, L. A. & Sankaran, V. G. Unraveling Hematopoiesis through the Lens of Genomics. Cell 182, 1384–1400 (2020).

25. Astle, W. J. et al. The Allelic Landscape of Human Blood Cell Trait Variation and Links to Common Complex Disease. Cell 167, 1415–1429.e19 (2016).

26. Notta, F. et al. Distinct routes of lineage development reshape the human blood hierarchy across ontogeny. Science 351, (2016).

27. Carrelha, J. et al. Hierarchically related lineage-restricted fates of multipotent haematopoietic stem cells. Nature 554, (2018).

28. Satpathy, A. T. et al. Massively parallel single-cell chromatin landscapes of human immune cell development and intratumoral T cell exhaustion. Nat. Biotechnol. 37, 925–936 (2019).

29. Mapping the human genetic architecture of COVID-19. Nature 600, 472–477 (2021).

30. Wakefield, J. A Bayesian measure of the probability of false discovery in genetic epidemiology studies. Am. J. Hum. Genet. 81, 208–227 (2007).

31. You, M. et al. Single-cell epigenomic landscape of peripheral immune cells reveals establishment of trained immunity in individuals convalescing from COVID-19. Nat. Cell Biol. 23, 620–630 (2021).

32. Wack, A. Monocyte and dendritic cell defects in COVID-19. Nature cell biology vol. 23 445–447 (2021).

33. Saichi, M. et al. Single-cell RNA sequencing of blood antigen-presenting cells in severe COVID-19 reveals multi-process defects in antiviral immunity. Nat. Cell Biol. 23, 538–551 (2021).

34. Arunachalam, P. S. et al. Systems biological assessment of immunity to mild versus severe COVID-19 infection in humans. Science 369, 1210–1220 (2020).

35. Pairo-Castineira, E. et al. Genetic mechanisms of critical illness in COVID-19. Nature 591, 92–98 (2021).

36. Kho, P. F. et al. Multi-tissue transcriptome-wide association study identifies eight candidate genes and tissue-specific gene expression underlying endometrial cancer susceptibility. Commun Biol 4, 1211 (2021).

37. Horowitz, J. E. et al. Genome-wide analysis in 756,646 individuals provides first genetic evidence that expression influences COVID-19 risk and yields genetic risk scores predictive of severe disease. medRxiv (2021) doi:10.1101/2020.12.14.20248176.

38. Schep, A. N., Wu, B., Buenrostro, J. D. & Greenleaf, W. J. chromVAR: inferring transcription-factor-associated accessibility from single-cell epigenomic data. Nat. Methods 14, 975–978 (2017).

39. Schmidl, C. et al. Transcription and enhancer profiling in human monocyte subsets. Blood 123, e90–9 (2014).

40. Reyes, M. et al. An immune-cell signature of bacterial sepsis. Nat. Med. 26, 333–340 (2020).

41. Rosenbauer, F. & Tenen, D. G. Transcription factors in myeloid development: balancing differentiation with transformation. Nat. Rev. Immunol. 7, 105–117 (2007).

42. Schulte-Schrepping, J. et al. Severe COVID-19 Is Marked by a Dysregulated Myeloid Cell Compartment. Cell 182, 1419–1440.e23 (2020).

43. Kachuri, L. et al. Genetic determinants of blood-cell traits influence susceptibility to childhood acute lymphoblastic leukemia. Am. J. Hum. Genet. 108, 1823–1835 (2021).

44. Jeon, S. et al. Genome-wide trans-ethnic meta-analysis identifies novel susceptibility loci for childhood acute lymphoblastic leukemia. Leukemia (2021) doi:10.1038/s41375-021-01465-1.

45. Cazzola, A. et al. Prenatal Origin of Pediatric Leukemia: Lessons From Hematopoietic Development. Front. Cell Dev. Biol. 0, (2021).

46. Fitch, B. et al. Human pediatric B-cell acute lymphoblastic leukemias can be classified as B-1 or B-2-like based on a minimal transcriptional signature. Exp. Hematol. 90, 65–71.e1 (2020).

47. Single-Cell Trajectory Detection Uncovers Progression and Regulatory Coordination in Human B Cell Development. Cell 157, 714–725 (2014).

48. Studd, J. B. et al. Genetic predisposition to B-cell acute lymphoblastic leukemia at 14q11.2 is mediated by a CEBPE promoter polymorphism. Leukemia 33, 1–14 (2018).

49. Wiemels, J. L. et al. A functional polymorphism in the CEBPE gene promoter influences acute lymphoblastic leukemia risk through interaction with the hematopoietic transcription factor Ikaros. Leukemia 30, 1194–1197 (2015).

50. Mullighan, C. G. et al. Genome-wide analysis of genetic alterations in acute lymphoblastic leukaemia. Nature 446, 758–764 (2007).

51. Shah, S. et al. A recurrent germline PAX5 mutation confers susceptibility to pre-B cell acute lymphoblastic leukemia. Nat. Genet. 45, 1226–1231 (2013).

52. Duployez, N. et al. Germline PAX5 mutation predisposes to familial B-cell precursor acute lymphoblastic leukemia. Blood 137, 1424–1428 (2021).

53. Bartosovic, M., Kabbe, M. & Castelo-Branco, G. Single-cell CUT&Tag profiles histone modifications and transcription factors in complex tissues. Nat. Biotechnol. 39, 825–835 (2021).

54. Wu, S. J. et al. Single-cell CUT&Tag analysis of chromatin modifications in differentiation and tumor progression. Nat. Biotechnol. 39, 819–824 (2021).

55. Pliner, H. A. et al. Cicero Predicts cis-Regulatory DNA Interactions from Single-Cell Chromatin Accessibility Data. Mol. Cell 71, 858–871.e8 (2018).

56. Fulco, C. P. et al. Activity-by-contact model of enhancer–promoter regulation from thousands of CRISPR perturbations. Nat. Genet. 51, 1664–1669 (2019).

57. Akgol Oksuz, B. et al. Systematic evaluation of chromosome conformation capture assays. Nat. Methods 18, 1046–1055 (2021).

58. Kaya-Okur, H. S. et al. CUT&Tag for efficient epigenomic profiling of small samples and single cells. Nat. Commun. 10, 1930 (2019).

59. Cusanovich, D. A. et al. Multiplex single cell profiling of chromatin accessibility by combinatorial cellular indexing. Science 348, 910–914 (2015).

60. Granja, J. M. et al. ArchR is a scalable software package for integrative single-cell chromatin accessibility analysis. Nat. Genet. 53, 403–411 (2021).

61. Stuart, T., Srivastava, A., Madad, S., Lareau, C. A. & Satija, R. Single-cell chromatin state analysis with Signac. Nat. Methods 18, 1333–1341 (2021).

62. Haghverdi, L., Lun, A. T. L., Morgan, M. D. & Marioni, J. C. Batch effects in single-cell RNA-sequencing data are corrected by matching mutual nearest neighbors. Nat. Biotechnol. 36, 421–427 (2018).

63. Tong, H., Faloutsos, C. & Pan, J.-Y. Fast random walk with restart and its applications. in Sixth International Conference on Data Mining (ICDM’06) (IEEE, 2006). doi:10.1109/icdm.2006.70.

64. McInnes, L., Healy, J. & Melville, J. UMAP: Uniform Manifold Approximation and Projection for Dimension Reduction. (2018).

65. Traag, V. A., Waltman, L. & van Eck, N. J. From Louvain to Leiden: guaranteeing well-connected communities. Sci. Rep. 9, 5233 (2019).

66. Yang, J., Lee, S. H., Goddard, M. E. & Visscher, P. M. GCTA: a tool for genome-wide complex trait analysis. Am. J. Hum. Genet. 88, 76–82 (2011).

67. Benner, C. et al. FINEMAP: efficient variable selection using summary data from genome-wide association studies. Bioinformatics 32, 1493–1501 (2016).

68. Corces, M. R. et al. Lineage-specific and single-cell chromatin accessibility charts human hematopoiesis and leukemia evolution. Nat. Genet. 48, 1193–1203 (2016).

69. Chen, X. et al. Joint single-cell DNA accessibility and protein epitope profiling reveals environmental regulation of epigenomic heterogeneity. Nat. Commun. 9, 4590 (2018).

70. Xiong, L. et al. SCALE method for single-cell ATAC-seq analysis via latent feature extraction. Nat. Commun. 10, 4576 (2019).

