## Supplementary figures for "Variant to function mapping at single-cell resolution through network propagation"

### Supplementary Fig. 1

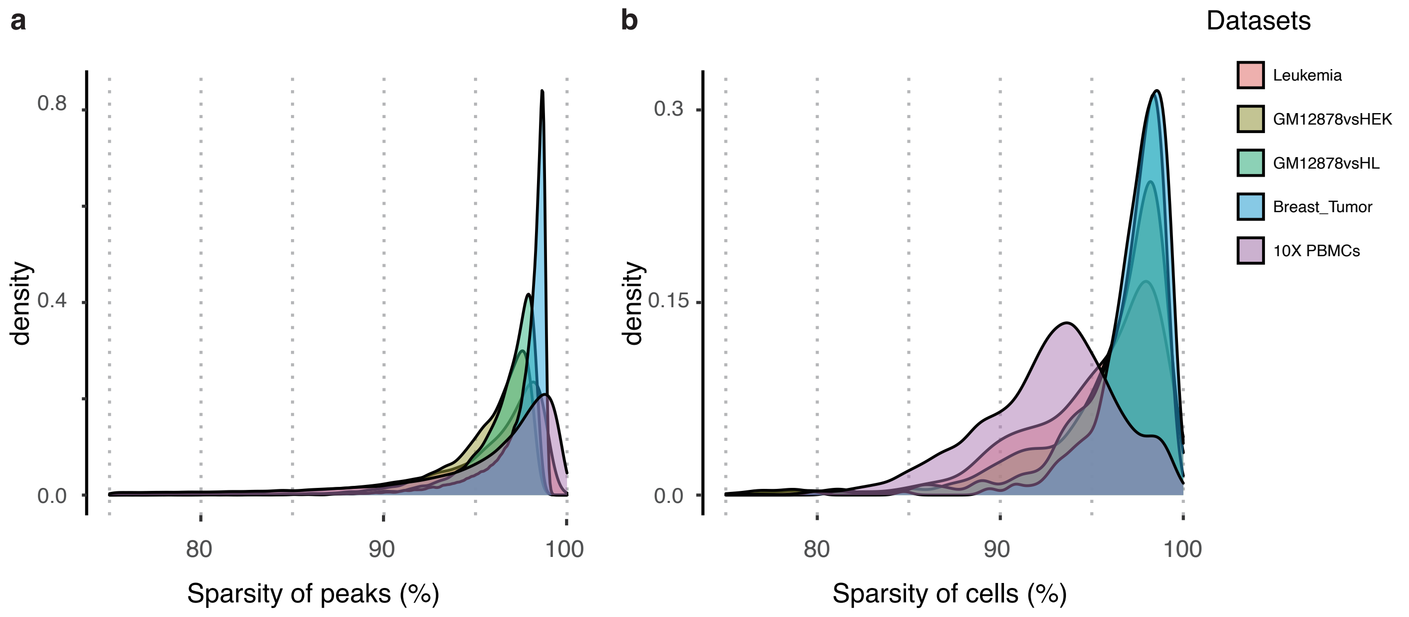


### Supplementary Fig. 2

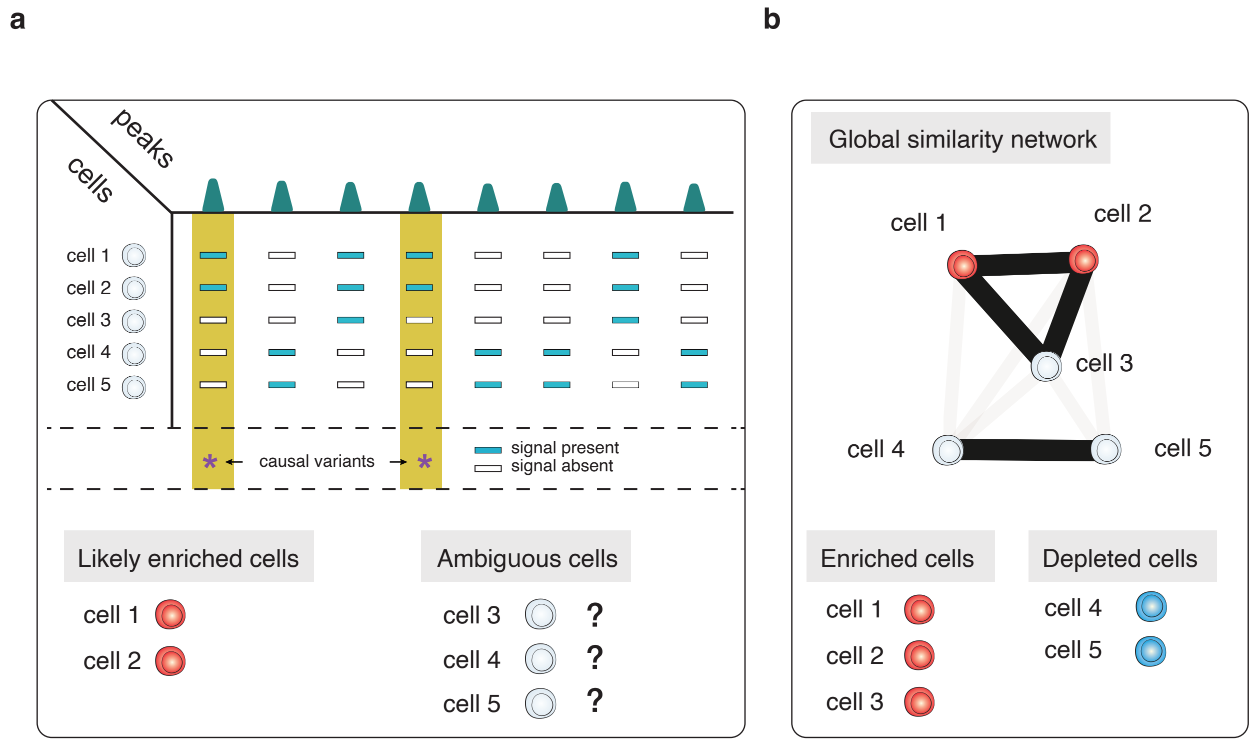

### Supplementary Fig. 3

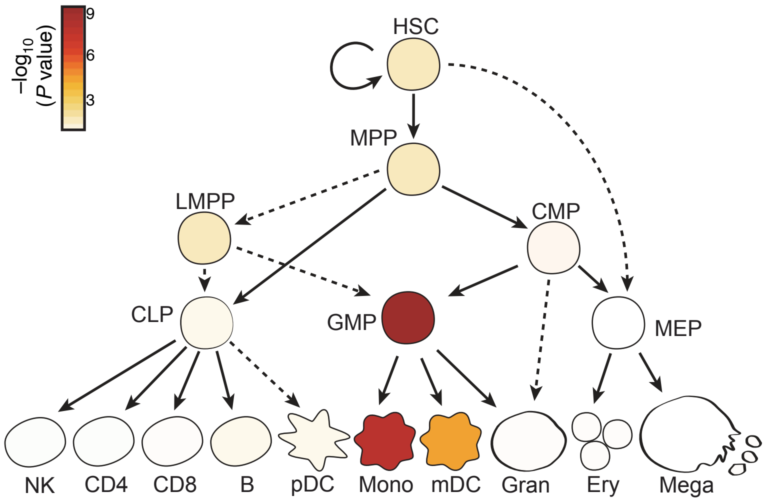


### Supplementary Fig. 4

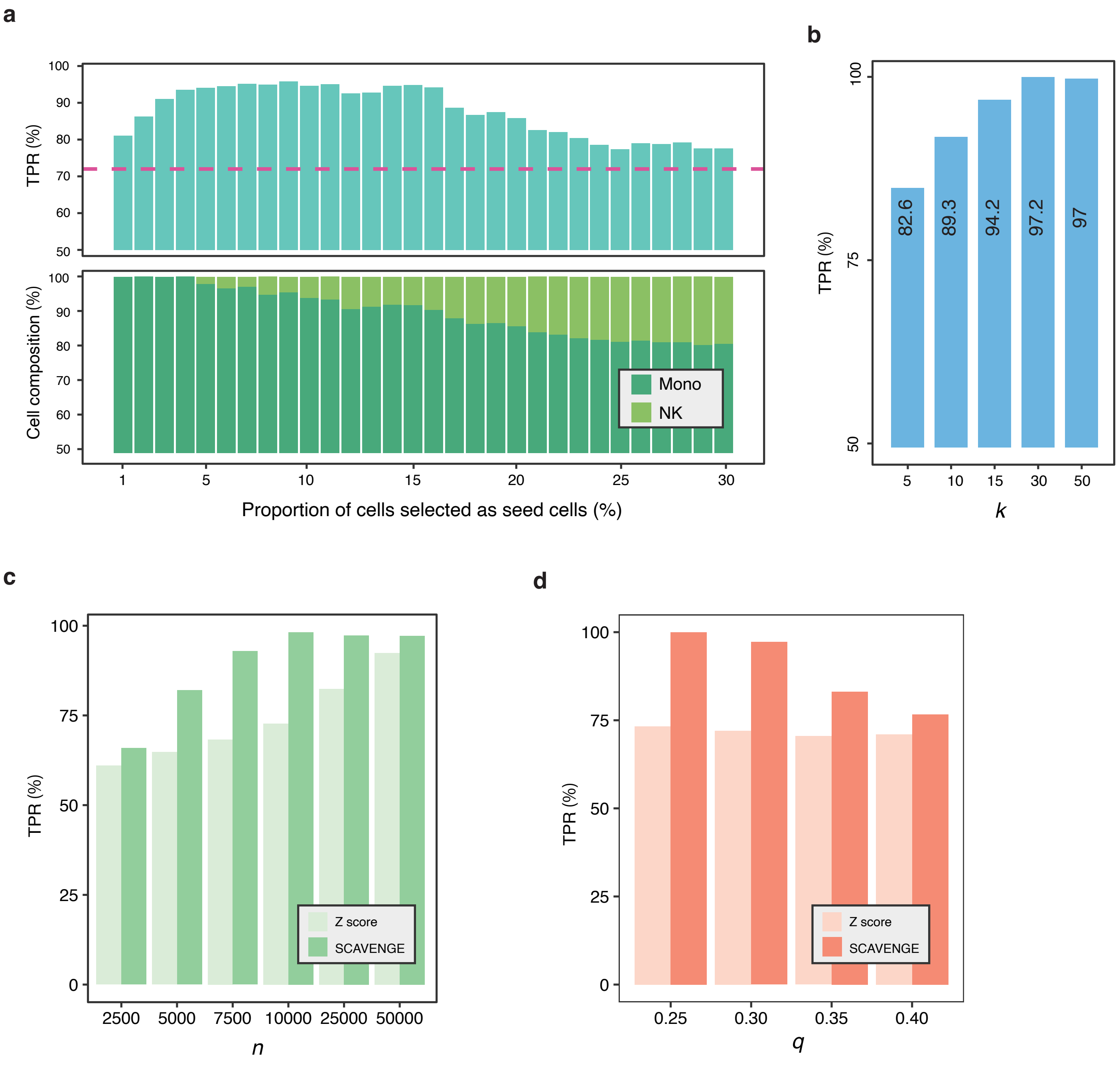

### Supplementary Fig. 5

**a**

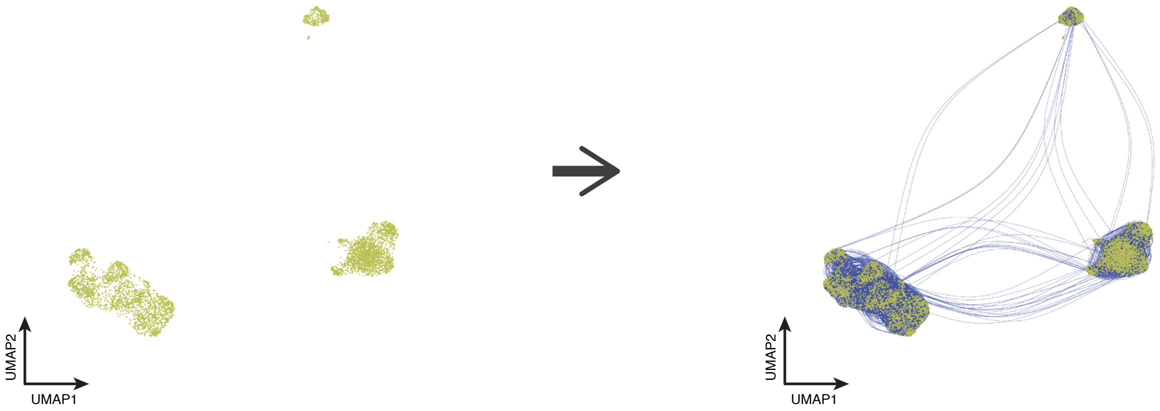


10X PBMC scATAC-seq (N=4,562)

**b**

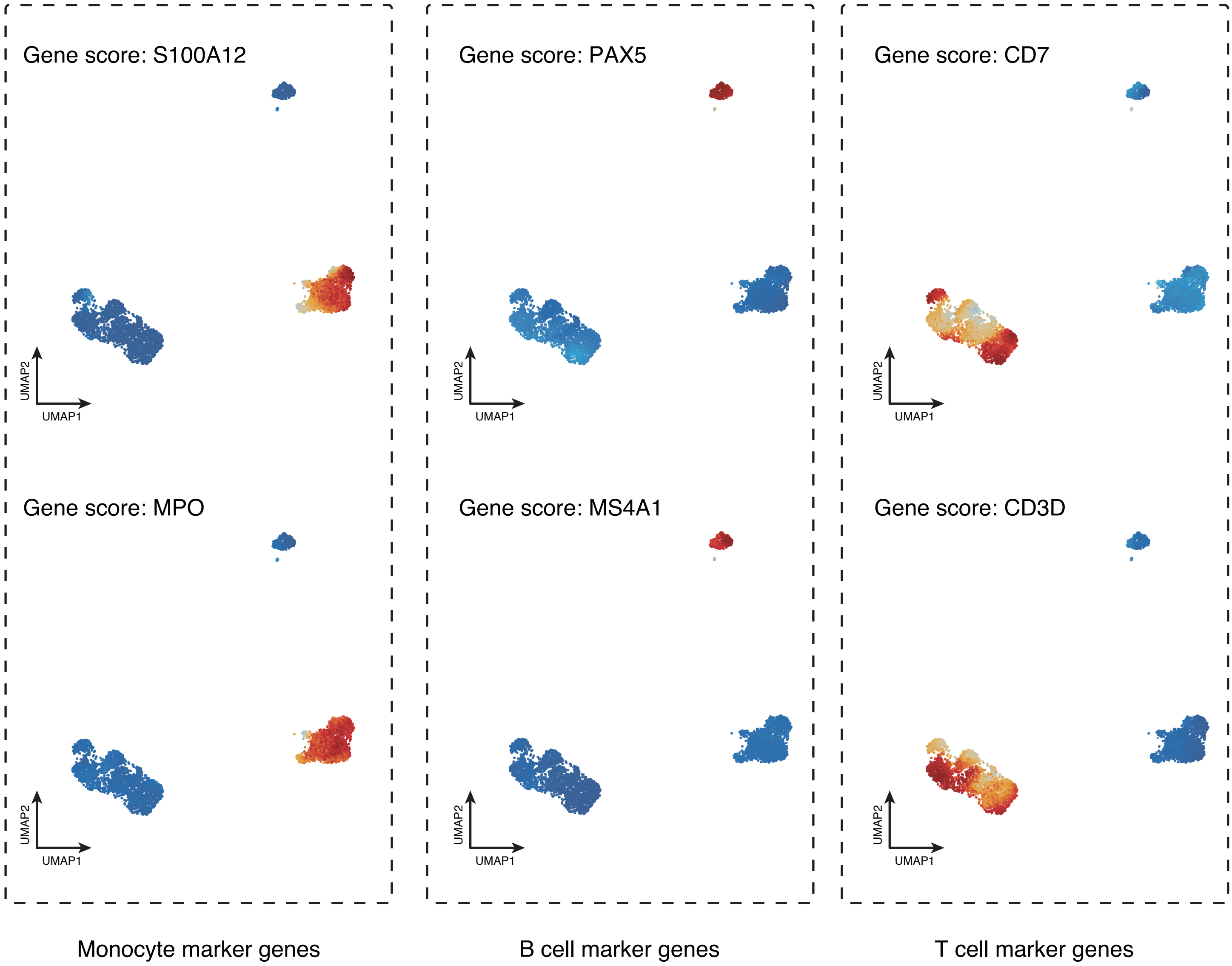

**Supplementary Fig. 6**

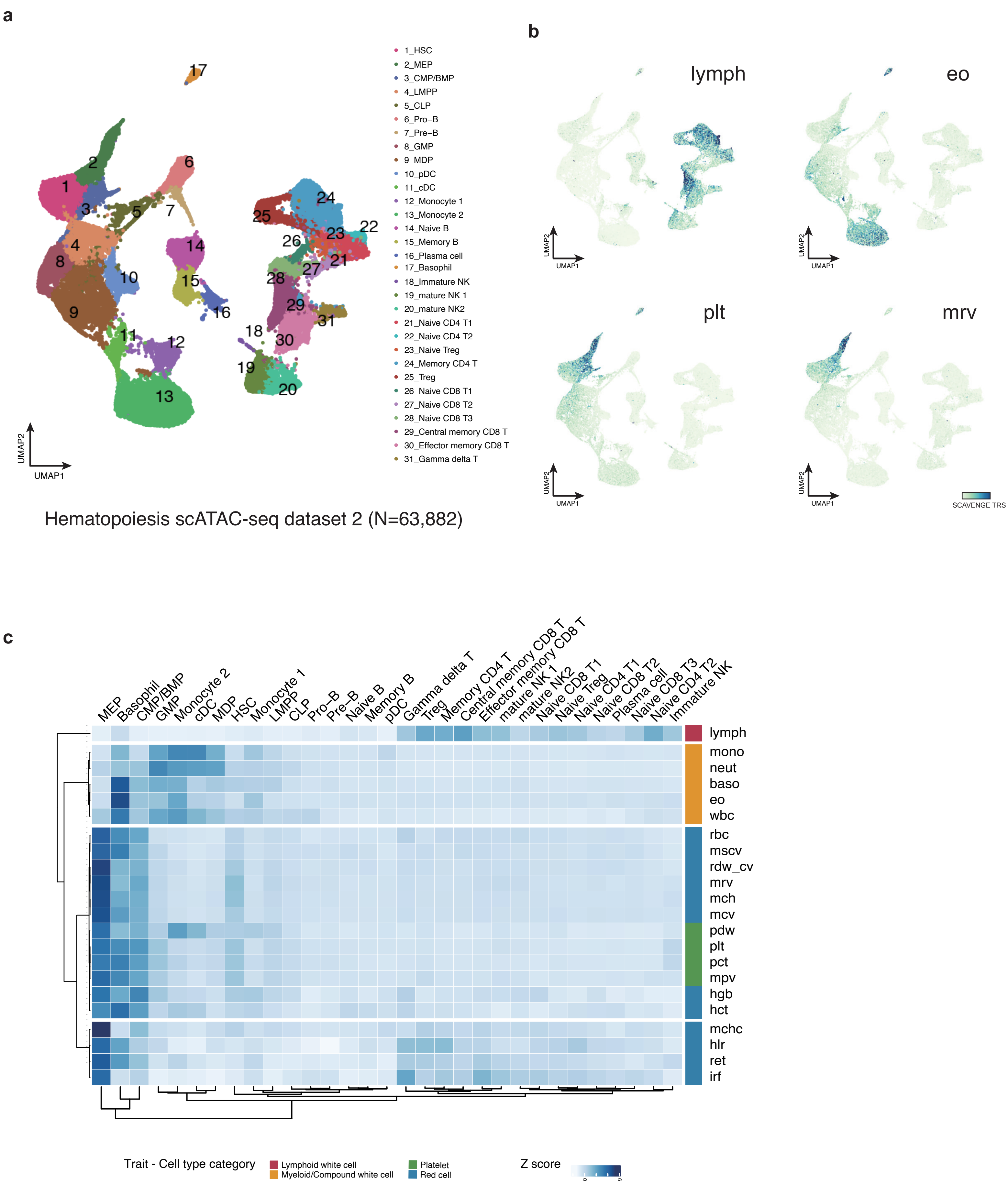

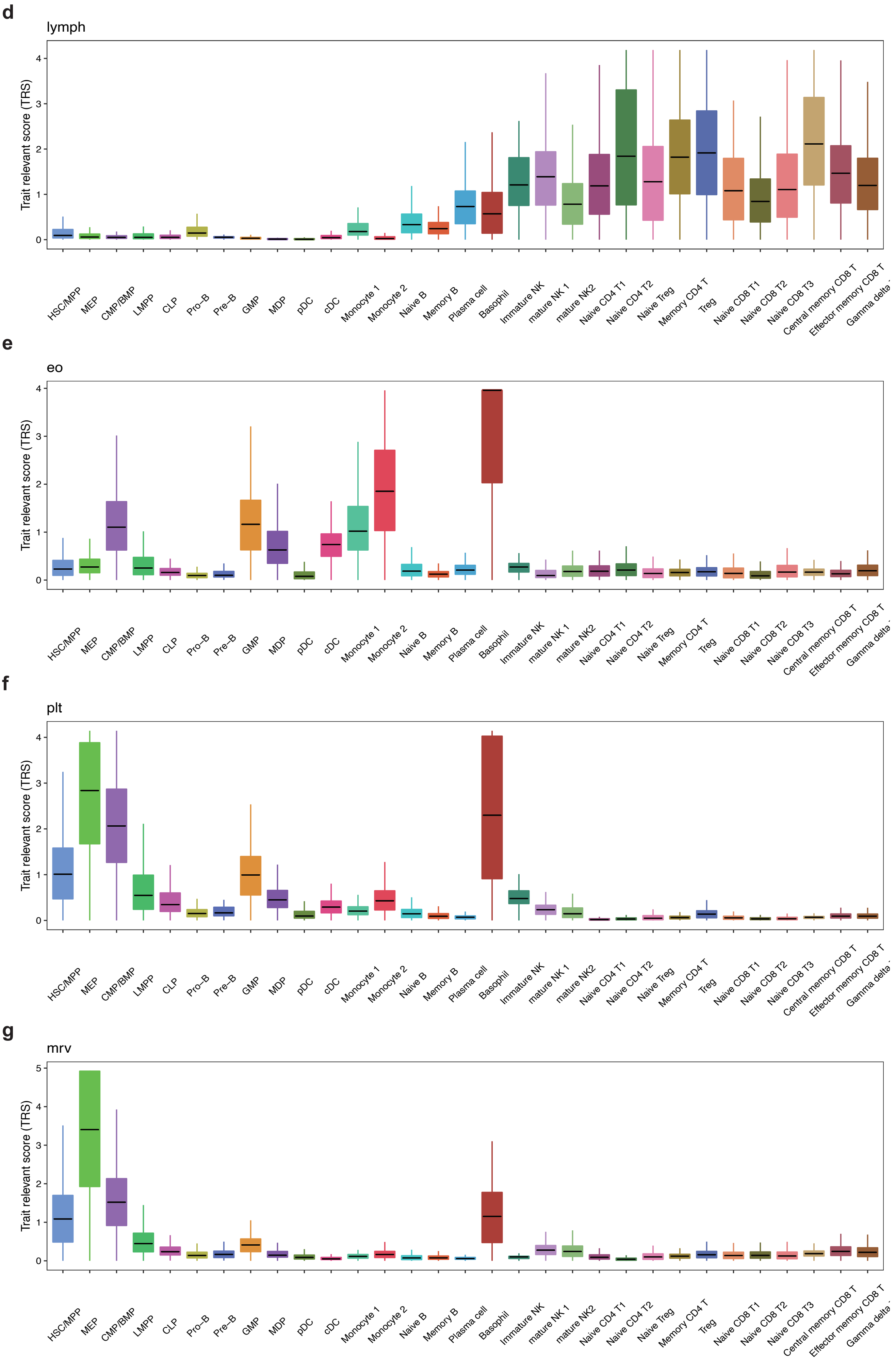
d
lymph
Trait relevant score (TRS)
e
eo
Trait relevant score (TRS)
f
plt
Trait relevant score (TRS)
g
mrv
Trait relevant score (TRS)
HSC/MPP
MEP
CMP/BMP
LMPP
CLP
Pro–B
Pre–B
GMP
MDP
pDC
cDC
Monocyte 1
Monocyte 2
Naive B
Memory B
Plasma cell
Basophil
Immature NK
mature NK 1
mature NK2
Naive CD4 T1
Naive CD4 T2
Naive Treg
Memory CD4 T
Treg
Naive CD8 T1
Naive CD8 T2
Naive CD8 T3
Central memory CD8 T
Effector memory CD8 T

### Supplementary Fig. 7

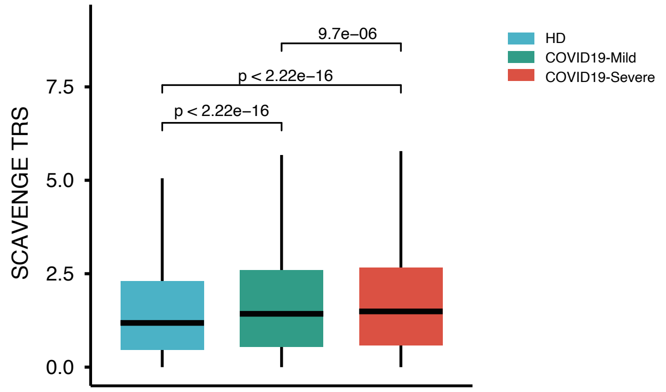


### Supplementary Fig. 8

**a**

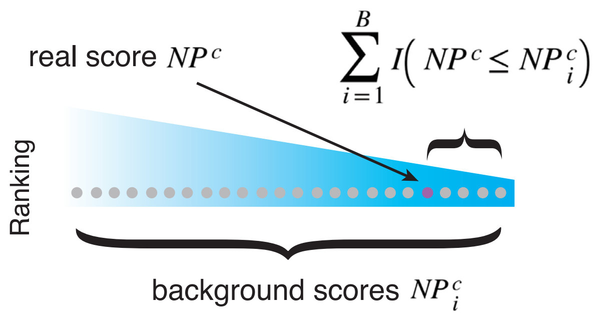


Empirical *P* value:

$$p_e^c = \frac{1 + \sum_{i=1}^{B} I(NP^c \leq NP_i^c)}{1 + B}$$

**b**

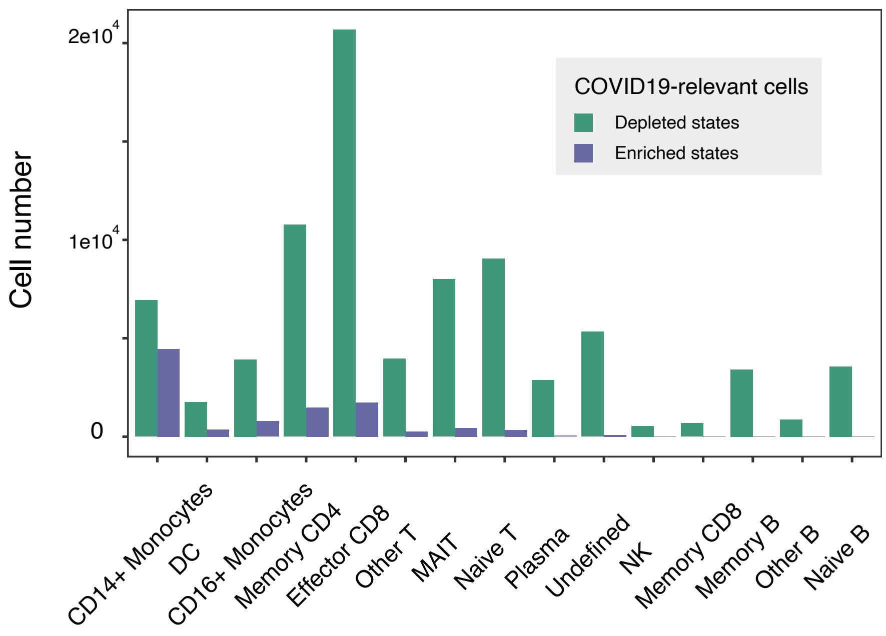

**Supplementary Fig. 9**

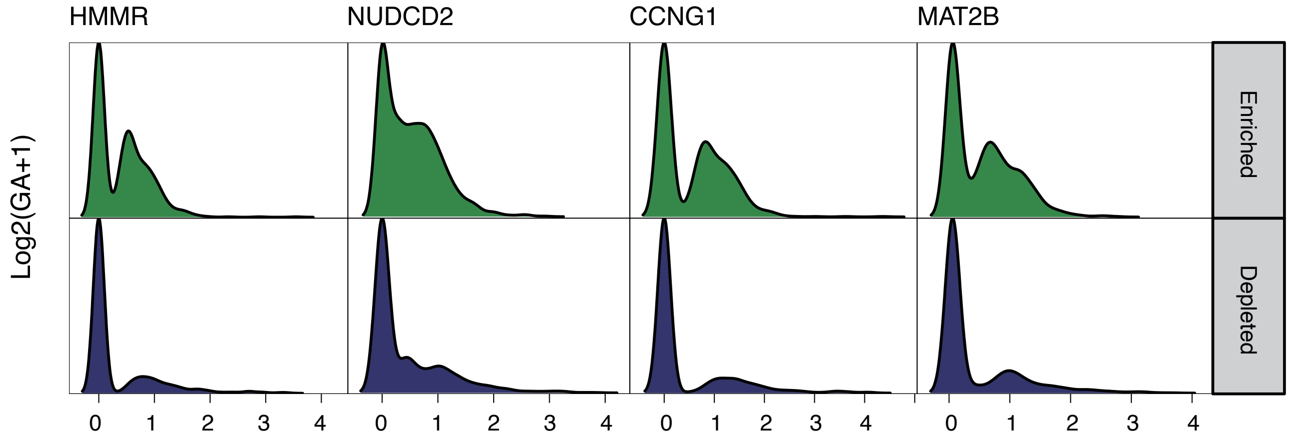


**Supplementary Fig. 10**

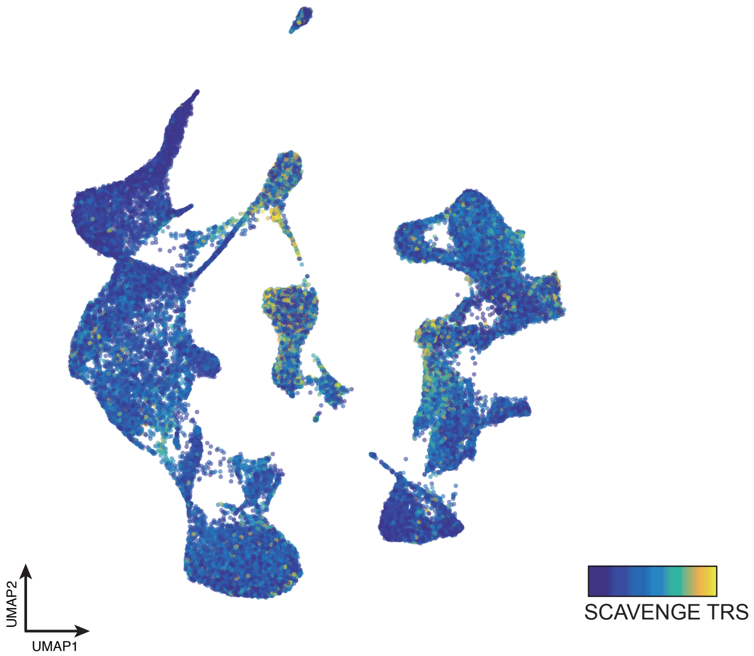


Hematopoiesis scATAC-seq dataset 2 (N=63,882)

### Supplementary Fig. 11

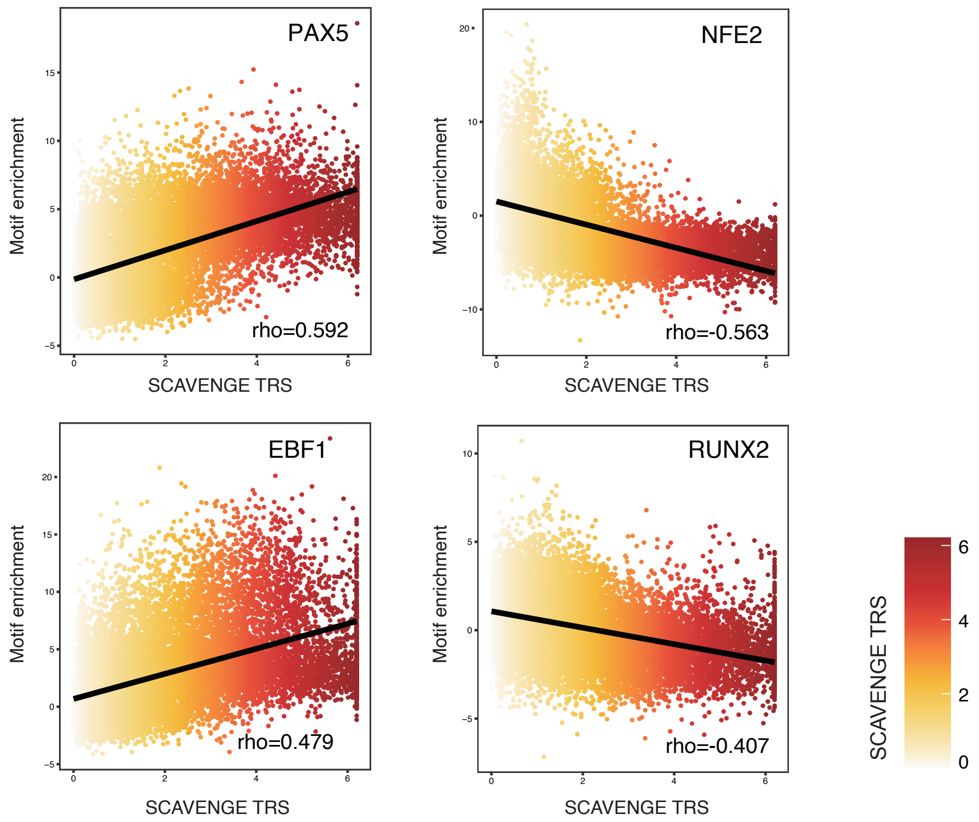
